# GLUCOCORTICOIDS REGULATE MITOCHONDRIAL FATTY ACID OXIDATION IN FETAL CARDIOMYOCYTES

**DOI:** 10.1101/2021.04.30.442128

**Authors:** Jessica R. Ivy, Roderic N. Carter, Jin-Feng Zhao, Charlotte Buckley, Helena Urquijo, Eva A. Rog-Zielinska, Emma Panting, Lenka Hrabalkova, Cara Nicholson, Emma J. Agnew, Matthew W. Kemp, Nicholas M. Morton, Sarah J. Stock, Caitlin Wyrwoll, Ian G. Ganley, Karen E. Chapman

**Affiliations:** University/BHF Centre for Cardiovascular Science, The Queen’s Medical Research Institute, The University of Edinburgh, 47 Little France Crescent, Edinburgh, EH16 4TJ, UK; Medical Research Council Protein Phosphorylation and Ubiquitylation Unit, University of Dundee, Dundee DD1 5EH; School of Human Sciences, The University of Western Australia, Crawley, WA6009, Australia; The Centre for Reproductive Health, The Queen’s Medical Research Institute, The University of Edinburgh, 47 Little France Crescent, Edinburgh, EH16 4TJ, UK; Department of Obstetrics and Gynaecology, Yong Loo Lin School of Medicine, National University of Singapore, 1E Kent Ridge Road, Singapore 119228, Republic of Singapore; Division of Obstetrics and Gynaecology, The University of Western Australia, Crawley, WA 6009, Australia; Centre for Perinatal and Neonatal Medicine, Tohoku University Hospital, Sendai, Japan; The Usher Institute, The University of Edinburgh, 47 Little France Crescent, Edinburgh, EH16 4UX, UK

**Author notes:** EJA: Food Standards Scotland, Q Spur, Saughton House, Broomhouse Dr, Edinburgh, EH11 3XD. ER-Z: Institute for Experimental Cardiovascular Medicine, University Heart Center Freiburg · Bad Krozingen, and Faculty of Medicine, University of Freiburg, Freiburg, Germany. CB: Strathclyde Institute of Pharmacy and Biomedical Sciences, 161 Cathedral Street, Glasgow, G4 0RE. CORRESPONDENCE: Karen E. Chapman, University/BHF Centre for Cardiovascular Science, The Queen’s Medical Research Institute, 47 Little France Crescent, Edinburgh, EH16 4TJ.

**Keywords:** Glucocorticoid, cardiomyocytes, early-life programming, heart, antenatal corticosteroids, preterm birth

## Abstract

The late gestational rise in glucocorticoids contributes to the structural and functional maturation of the perinatal heart. Here, we hypothesised that glucocorticoid action contributes to the metabolic switch in perinatal cardiomyocytes from carbohydrate to fatty acid oxidation. In primary mouse fetal cardiomyocytes, dexamethasone treatment induced expression of genes involved in fatty acid oxidation and increased mitochondrial oxidation of palmitate, dependent upon glucocorticoid receptor (GR). Dexamethasone did not, however, induce mitophagy or alter the morphology of the mitochondrial network. In neonatal mice, dexamethasone treatment induced cardiac expression of fatty acid oxidation genes *in vivo.* However, dexamethasone treatment of pregnant C57Bl/6 mice at embryonic day (E)13.5 or E16.5 failed to induce fatty acid oxidation genes in fetal hearts assessed 24 hours later. Instead, at E17.5, fatty acid oxidation genes were down-regulated by dexamethasone, as was GR itself. PGC-1α, required for glucocorticoid-induced maturation of primary mouse fetal cardiomyocytes *in vitro*, was down-regulated *in vivo* in fetal hearts at E17.5, 24 hours after dexamethasone administration. Similarly, following a course of antenatal corticosteroids in a sheep model of preterm birth, both GR and PGC-1α were down-regulated in fetal heart. These data suggest endogenous glucocorticoids support the perinatal switch to fatty acid oxidation in cardiomyocytes through changes in gene expression rather than gross changes in mitochondrial volume or mitochondrial turnover. Moreover, our data suggest that treatment with exogenous glucocorticoids may interfere with normal fetal heart maturation, possibly by down-regulating GR. This has implications for clinical use of antenatal corticosteroids when preterm birth is considered a possibility.

## INTRODUCTION

The dramatic increase in fetal glucocorticoid hormone concentration in late gestation is essential to support the transition from intrauterine to extrauterine life (Hillman *et al.*, 2012; Rog-Zielinska *et al.*, 2014). Administration of synthetic corticosteroids (betamethasone or dexamethasone) to pregnant women at risk of preterm delivery is standard care in high and middle-income countries, with the aim of maturing the fetus to reduce neonatal morbidity and mortality (Kemp *et al.*, 2016; Agnew *et al.*, 2018). In addition to the well-known effects on lung maturation (Cole *et al.*, 1995; Bird *et al.*, 2015; Laresgoiti *et al.*, 2016), glucocorticoids promote pro-survival adaptions in neonatal energy metabolism and in the cardiovascular system (Hillman *et al.*, 2012). However, which of these effects are directly attributable to glucocorticoid activation of GR within tissues and which are mediated by other factors remains uncertain. Also unclear is whether antenatal administration of synthetic glucocorticoids mimics endogenous glucocorticoid action in the fetal cardiovascular system. Our previous data suggest antenatal dexamethasone treatment dysregulates cardiac function and down-regulates endogenous glucocorticoid action in the fetal heart (Agnew *et al.*, 2019), potentially altering the normal trajectory of perinatal cardiac maturation.

The normal increase in fetal glucocorticoids in late gestation supports neonatal blood pressure (Hillman *et al.*, 2012) and is essential to structurally and functionally mature the fetal heart (Rog-Zielinska *et al.*, 2013). *In utero*, our 'SMGRKO' mice, with *Sm22-Cre*-mediated GR deficiency in cardiomyocytes and vascular smooth muscle cells show impaired heart function, disrupted cardiac ultrastructure and fail to induce key genes required for cardiac contractile function, calcium handling and energy metabolism (Rog-Zielinska *et al.*, 2013). Supporting direct effects of GR, glucocorticoid treatment of primary mouse fetal cardiomyocytes *in vitro* matures ultrastructure and increases contractile function, mitochondrial capacity (O_2_ consumption rate, basally and after uncoupling of mitochondria) and markers of cardiomyocyte maturation (Rog-Zielinska *et al.*, 2015). Similarly, in human embryonic stem cell (ESC)-derived cardiomyocytes treated with dexamethasone, contractile force is increased and systolic calcium transient decay is faster (Kosmidis *et al.*, 2015).

During the transition to a higher oxygen environment and a greater cardiac workload at birth, the cardiac preference for energy substrate switches. The fetal heart derives most of its ATP from glucose and lactate oxidation, with only a minor contribution from fatty acids. After birth, the increased demand for ATP is met primarily by oxidation of long chain fatty acids (Lopaschuk & Jaswal, 2010). This is associated with increased mitochondrial functional capacity. PGC-1α, a master transcriptional regulator of mitochondrial capacity, is expressed in the late gestation fetal heart and expression increases markedly after birth (Lehman *et al.*, 2000). Mice with global knock-out of PGC-1α show 50% mortality before weaning (Lin *et al.*, 2004), suggesting it is important in the perinatal period. PGC-1α is a glucocorticoid target gene and, *in vivo* in fetal heart, is induced 6 hours after glucocorticoid treatment (Rog-Zielinska *et al.*, 2015). PGC-1α is also induced *in vitro* in primary mouse fetal cardiomyocytes (Rog-Zielinska *et al.*, 2015). Here, the GR-mediated increase in PGC-1α expression is crucial for the glucocorticoid-induced maturation of myofibril structure and increased mitochondrial O_2_ consumption (Rog-Zielinska *et al.*, 2015). Knock-down of PGC-1α abolished both. RNAseq analysis performed on primary mouse fetal cardiomyocytes harvested 2 hours after glucocorticoid addition in the presence of cycloheximide (to block secondary effects) identified a number of differentially expressed genes, likely to be primary targets of GR (Rog-Zielinska *et al.*, 2015). As well as *Ppargc1a* (encoding PGC-1α), master regulators of mitochondrial fatty acid oxidation (*Klf15, Lipin1, Cebpb, Ppara*) were induced. This suggests that glucocorticoid action may promote the perinatal switch in cardiomyocytes from carbohydrate to fatty acid oxidation as the preferred substrate for ATP generation.

Cellular differentiation is often associated with metabolic remodelling and the autophagic turnover of mitochondria by mitophagy (Rodger *et al.*, 2018). Mitophagic removal of small fetal mitochondria in perinatal cardiomyocytes is reportedly a prerequisite for the formation of morphologically distinct adult mitochondria and maturation into cardiomyocytes optimised for fatty acid metabolism (Gong *et al.*, 2015). The triggers for mitophagy in perinatal cardiomyocytes are currently unknown. *mito*-QC transgenic mice have a pH-sensitive fluorescent mitochondrial signal that monitors mitophagy *in vivo* (McWilliams *et al.*, 2016). These mice have revealed that mitophagy is occurring at E17.5 in the mouse fetal heart (McWilliams *et al.*, 2016), a time co-incident with peak GR activation in heart (Rog-Zielinska *et al.*, 2013). Furthermore, *Bnip3*, implicated in mitophagy, is a direct GR target gene in primary fetal cardiomyocytes (Rog-Zielinska *et al.*, 2015), raising the possibility that glucocorticoids may be a trigger for mitophagy in perinatal cardiomyocytes.

Here, we hypothesised that glucocorticoids increase fetal cardiomyocyte capacity for fatty acid oxidation. We also asked if any glucocorticoid-mediated increase in mitochondrial fatty acid oxidation capacity involves mitochondrial remodelling by mitophagy.

## MATERIALS AND METHODS

### Animals

Experiments involving mice were approved by the University of Edinburgh Animal Welfare and Ethical Review Body and carried out in strict accordance with accepted standards of humane animal care under the auspices of the Animal (Scientific Procedures) Act UK 1986. Mice were maintained under controlled lighting and temperature. C57BL/6J/Ola/Hsd (C57Bl/6J) mice were purchased from Harlan, then bred in house. *GR^+/−^* mice, heterozygous for a null mutation in the *Nr3c1* gene encoding GR (*Nr3c1^gtESK92MRCHGU^* mice), have been previously described (Michailidou *et al.*, 2008; Rog-Zielinska *et al.*, 2013). *GR^+/−^* mice, congenic on the C57Bl/6J background (>12 generations) were intercrossed to give *GR^+/+^*, *GR^+/−^* and *GR^−/−^* fetal littermates. The morning of the day the vaginal plug was found was designated E0.5. Fetuses were collected at E17.5, hearts were dissected and rapidly frozen on dry ice. Genotyping of fetal tissue by PCR used *LacZ* primers for the *Nr3c1^gtESK92MRCHGU^* (5’-GAGTTGCGTGACTACCTACGG-3’ and 5’-GTACCACAGCGGATGGTTCGG-3’) and wild-type *GR* alleles as described (Michailidou *et al.*, 2008). *mito*-QC mice, also on a C57Bl/6J background, have been described (McWilliams *et al.*, 2016). *mito*-QC heterozygous fetuses were used for fetal cardiomyocyte cultures. Fetuses were collected and cardiomyocytes isolated as described below.

For dexamethasone treatment, pregnant C57Bl/6J females (time-mated with C57Bl/6J males) were semi-randomised to experimental group (alternating groups as lifted from the cage) and injected (~0.1ml, intra-peritoneal) with dexamethasone (0.5mg/kg; Sigma-Aldrich, Poole, UK) or vehicle (5% ethanol) at either E13.5 or E16.5 and euthanised 24 hours later (E14.5 and E17.5, respectively). Fetuses were removed to ice-cold PBS. Hearts were excised and frozen on dry ice. Pregnant *GR^+/−^* dams were euthanised at E17.5 and fetal hearts removed and frozen as above. Neonatal C57Bl/6J mice were injected (intra-peritoneal) with dexamethasone (0.5mg/kg) or vehicle (5% ethanol) at post-natal day 1 (P1; day of birth being P0) and euthanised by decapitation 24 hours later. Hearts were removed and frozen as above. Tissues were identified by animal ID (blinding to genotype/treatment group) and stored at −80°C prior to analysis.

Sheep protocols were approved by the animal ethics committee of The University of Western Australia (RA/3/100/1452). Date-mated merino ewes carrying singleton pregnancies were randomized to receive 2 injections (intra-muscular) spaced by 24 hours of either saline (control) or betamethasone acetate with betamethasone phosphate (Celestone Chronodose, Merck & Co., Inc, Kenilworth, NJ) 0.25mg/kg per injection. A third group received a single injection of betamethasone acetate (0.125mg/kg). Betamethasone acetate was a gift from Merck & Co. as a preparation of betamethasone acetate equivalent to that in Celestone. Merck & Co. did not participate in the design, execution, or analysis of the study. The 0.25mg/kg Celestone dose approximates the clinical dose of 12mg of betamethasone for a 50kg woman and was the same dose used for our previous studies (Kemp *et al.*, 2018; Schmidt *et al.*, 2019). To reduce the risk of preterm labour from antenatal corticosteroids, all animals, irrespective of subsequent treatment, received one intramuscular dose of 150mg medroxyprogesterone acetate (Depo-Provera, Pfizer, New York, NY), five days before corticosteroid treatment. No other doses of medroxyprogesterone acetate were administered, nor were other tocolytics administered. All animals were delivered at 122±1 days (term being ~147 days).

Two days after their initial steroid or saline treatment, ewes received an intravenous injection of ketamine (10mg/kg) and midazolam (0.5mg/kg). A spinal injection of 3ml lignocaine (20mg/ml) was then administered and surgical delivery commenced. The lamb received an intramuscular injection of ketamine (10mg/kg) before placing a 4.5mm endotracheal tube by tracheostomy. Lambs were weighed, dried, and placed in an infant warmer (Fisher & Paykel Healthcare, New Zealand). Intermittent positive pressure ventilation was performed using Acutronic Fabian infant ventilators (Acutronic Medical System, Hirzel, Switzerland) and maintained for 30 minutes using the following settings: initial peak inspiratory pressure (PIP) of 40 cmH_2_O, positive end expiratory pressure (PEEP) of 5 cmH_2_O, respiratory rate of 50 breaths per minutes, inspiratory time of 0.5 seconds. Gas mix was 100% heated and humidified oxygen. PIP was titrated to achieve a tidal volume of 7ml/kg. Lambs were euthanised with an IV overdose of pentobarbitone sodium at 160mg/kg. Necropsy was performed immediately following a 30 minute ventilation procedure which commenced at delivery. At necropsy (within 40 minutes of delivery), left and right ventricles of the heart were dissected and samples snap frozen.

### Fetal Cardiomyocyte Cultures

Primary fetal cardiomyocyte cultures were prepared as described (Rog-Zielinska *et al.*, 2015). Briefly, hearts were rapidly dissected from 5-30 E14.5-E15.5 C57Bl/6J or *mito*-QC fetuses, placed in warm Tyrode’s salt solution containing 0.1% sodium bicarbonate then rinsed in complete medium (DMEM supplemented with 100IU/ml penicillin, 100mg/ml streptomycin, 10% fetal bovine serum, 0.1% non-essential amino acids, Sigma-Aldrich, Poole, UK) before digestion at 37oC with gentle agitation for 10 minutes in 5ml enzyme buffer (PBS supplemented with 0.8% NaCl, 0.2% D-(+)-glucose, 0.02% KCl, 0.000575% NaH_2_PO_4_·H_2_O, 0.1% NaHCO_3_, pH7.4) containing 0.03% type II collagenase (Worthington Biochemical Corp, Lakewood, New Jersey, USA), 0.125% porcine pancreatin (Sigma-Aldrich, Poole, UK). After 10 minutes, isolated cells and enzyme buffer were removed and the enzymatic reaction quenched by adding the same volume of complete medium. Fresh enzyme buffer was then added to the hearts. Approximately 8 digestions were performed on the hearts until their structure was lost. Isolated cells were centrifuged at 1000 rpm for 10 minutes at room temperature, the supernatant removed and pellets pooled. Pooled cells were centrifuged again and resuspended in 15ml isolation buffer (Ham’s F12 supplemented with 100IU/ml penicillin, 100mg/ml streptomycin, 0.002% ascorbic acid, 1% fetal bovine serum, 0.1176% NaHCO_3_). To reduce the number of fibroblasts, cells were incubated in a tissue culture plate (ThermoFisher, UK) at 37°C, 5% CO_2_ for 3 hours during which fibroblasts adhered to the plastic. Non-adherent cells were aspirated, centrifuged, resuspended in 1ml of complete medium and seeded at a density of 0.25 × 10^6^cells/ml for extra-cellular flux (ECF; Seahorse) assays, RNA analysis, or mitochondrial morphology. This protocol yields ≥98% cardiomyocytes (troponin T+ cells) (Rog-Zielinska *et al.*, 2015). Spontaneous beating of cardiomyocytes was observed within 12 hours. For ECF assays, cardiomyocytes were seeded onto 24 well gelatin (Sigma, UK) coated V7 Seahorse plates (Agilent Technologies LDA UK Ltd, Stockport, Cheshire, UK) in complete medium then treated with dexamethasone (1μM) or vehicle (0.01% ethanol) for 24 hours prior to ECF assay (below). We have previously shown that this dose of dexamethasone elicits maximal glucocorticoid responses in fetal cardiomyocytes (Rog-Zielinska *et al.*, 2015). For RNA analysis, primary fetal cardiomyocytes were seeded in 12-well gelatin-coated tissue culture plates for 48 hours. Cells were treated with dexamethasone (1μM) or vehicle (0.01% ethanol) and lysed 6 or 24 hours later by adding 0.5ml TRIzol (Invitrogen, ThermoFisher, UK) following removal of medium. For measurement of mitochondrial morphology and mitophagy, cardiomyocytes were cultured on gelatin-coated glass chambered slides (Ibidi μ-Slide 4 well Glass bottom, Thistle Scientific LTD, Glasgow, UK) prior to staining.

### Extra-cellular Flux (ECF) Assays

ECF assays were carried out using a Seahorse XF^e^24 Bioanalyzer (Agilent Technologies LDA UK Ltd, Stockport, Cheshire, UK). All Seahorse reagents were purchased from Agilent Technologies LDA UK Ltd. All findings were reproduced in at least 2 experiments. For each experiment, treatment groups were randomised across the plate. Each well was assigned a number (based on the number of treatment groups) so that numbers were spread across the plate. Treatments were then randomised to numbers. On the day of the assay, complete culture medium was gently aspirated from the cells and exchanged for Seahorse assay medium (with supplements defined below). The cells were gently rinsed 3 times with Seahorse assay medium, leaving a final volume of 525μl for the assay. During all assays, 3 measurements at 2.5 minute intervals, were recorded at baseline and after each drug addition.

### Normalisation of Extra-cellular Flux Assays using sulforhodamine B (SRB) Assay

ECF assays were normalised to protein measured using sulforhodamine B (SRB) dye-based protein assay (Skehan *et al.*, 1990). Initial experiments confirmed the linearity of the SRB assay for use in quantifying primary cardiomyocyte protein levels (Supplementary Figure 1A). Following ECF assays, cells were fixed by addition of 50μl cold 50% trichloroacetic acid (Sigma-Aldrich, Poole, UK) per well and stored up to 1 week at 4^°^);C. Cells were then washed 10 times with tap water and air-dried. 50μl SRB solution (0.4% w/v sulforhodamine B dye (Sigma-Aldrich, Poole, UK) in 1% acetic acid (Sigma-Aldrich, Poole, UK)) was added to dried cells and incubated for 30 minutes at room temperature. Cells were then washed 4 times with 1% acetic acid and air-dried. Cell-bound dye was re-dissolved in 200μl 10mM Tris pH 10.5. Absorbance was measured at a wavelength of 540nm.

**Figure 1.**
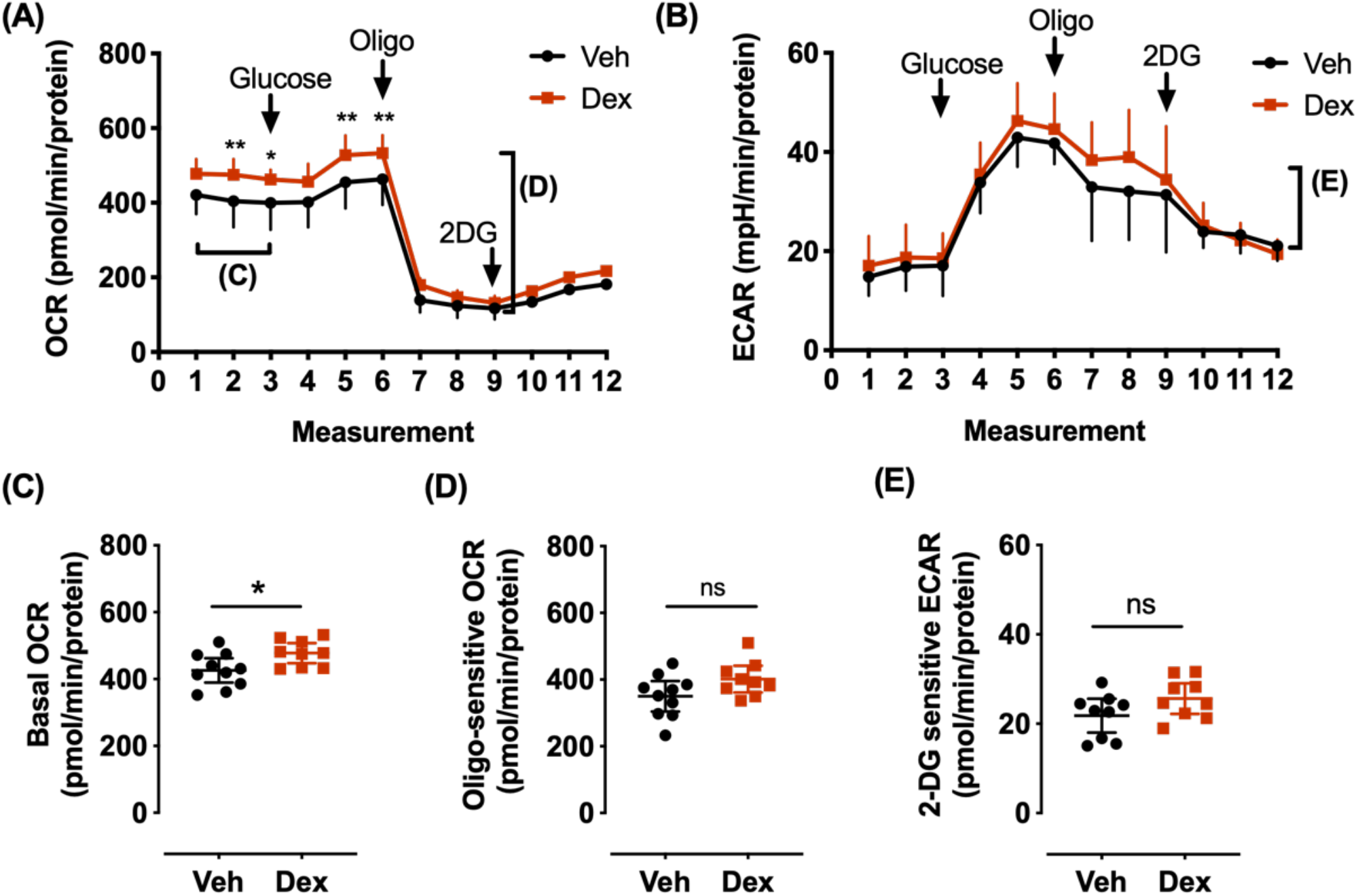
Dexamethasone increased basal respiration in primary fetal cardiomyocytes. Primary fetal cardiomyocytes were prepared by digesting pooled E14.5-15.5 C57BL/6J fetal hearts, then cultured for 48 hours in complete DMEM culture medium. Cardiomyocytes were treated with dexamethasone (Dex, 1μM, red) or vehicle (Veh, black). After 24 hours, medium was exchanged for Seahorse base medium and cardiomyocyte metabolism was analyzed by extra-cellular flux assay. After 3 basal measurements, glucose (10mM) was added, followed by oligomycin (Oligo, 1.5μM) and 2-deoxyglucose (2DG, 100mM). Oxygen consumption rate (OCR, **A**) and extra-cellular acidification rate (ECAR, **B**) were measured three times over 7.5 minutes following each addition. (**C**) Basal OCR was calculated as the mean of the 3 basal measurements. (**D**) ATP production was estimated as the maximum change in OCR following the addition of oligomycin (oligomycin-sensitive OCR). (**E**) Glycolysis (2DG-sensitive ECAR) was measured as the maximum change in ECAR folllowing addition of 2DG. Representative data from an experiment using cardiomyocytes pooled from tens of fetuses across 9-10 wells: data are mean ± SD, ns=not significant, **p<0.01 *p<0.05 (t-tests) (**B**), or two-way ANOVA followed by *post hoc* Sidak’s tests (**A**).

### Glycolysis Assays

Assays to estimate the rate of glycolysis in fetal cardiomyocytes were performed in two different ways. In the first, complete culture medium was exchanged for pre-warmed Seahorse Assay medium supplemented with 10mM glucose (Sigma-Aldrich, Poole, UK), 1mM sodium pyruvate (Sigma-Aldrich, Poole, UK). After basal OCR/ECAR measurements, 2-deoxyglucose (2DG; Sigma-Aldrich, Poole, UK, 100 mM) was added to inhibit glycolysis, followed by antimycin and rotenone (AR, Sigma-Aldrich, Poole, UK, 2μM) to inhibit respiration. The second measure was a Glycolysis Stress Test, performed according to the manufacturer’s protocol. Briefly, the cells were incubated in pre-warmed glucose-free Seahorse-XF Base medium for 1 hour in a (non-CO_2_) 37°C incubator prior to the assay. After 3 basal measurements, glucose (Sigma-Aldrich, Poole, UK, 10mM) was added to enable glycolysis, followed by oligomycin (Sigma-Aldrich, Poole, UK, 1.5μM) to inhibit respiratory ATP production and finally 2DG (Sigma-Aldrich, Poole, UK, 100 mM) to inhibit glycolysis.

### Fatty Acid Oxidation Assay

To test the ability of primary fetal cardiomyocytes to utilize long chain fatty acids, cells were pre-treated with etomoxir (Sigma-Aldrich, Poole, UK) to inhibit the CPT-1 mitochondrial fatty-acid uptake transporter, prior to performing a standard Seahorse Mitochondrial Stress Test. Initial experiments with high concentrations of etomoxir (40-160μM) reduced OCR in the presence of BSA-palmitate (Supplementary Figure 1C). However, within this range (120μM), etomoxir also showed likely off-target inhibition of mitochondrial respiration (Supplementary Figure 1B). This is consistent with an emerging literature showing that etomoxir can inhibit complex I at commonly used high doses (Divakaruni *et al.*, 2016; Yao *et al.*, 2018). Accordingly, we used 6μM etomoxir for all further experiments, a dose which inhibits fatty acid oxidation and avoids the CPT-1-independent effects associated with higher doses (Spurway *et al.*, 1997). Briefly, culture medium was exchanged for Seahorse Assay medium supplemented with 5mM glucose (Sigma-Aldrich, Poole, UK), 0.5mM carnitine (Sigma-Aldrich, Poole, UK). Cells were pre-treated with etomoxir (6μM in medium) or vehicle (medium) 15 minutes prior to the addition of BSA-Palmitate (100μM; Agilent Technologies LDA UK Ltd). The Seahorse Mitochondrial Stress test assay was started 15 minutes after the addition of BSA-Palmitate. Briefly, the test progressed as follows: basal respiration measurements, addition of oligomycin (1.5μM), addition of carbonyl cyanide-4-(trifluoromethoxy)phenylhydrazone (FCCP; 1μM), addition of AR (2μM). Non-mitochondrial respiration was calculated as the minimum OCR remaining following AR treatment. The mean of the 3 baseline measurements was used for the Basal OCR. Basal respiration was calculated by subtracting non-mitochondrial respiration from Basal OCR. ATP production was calculated as the maximum change in OCR following the addition of oligomycin and maximum respiration was calculated as the maximum OCR measurement induced by FCCP corrected for non-mitochondrial respiration. Leak respiration was calculated as the average oligomycin-insensitive OCR corrected for non-mitochondrial respiration.

### Measurements of Mitochondrial Morphology

On the day of staining of primary fetal cardiomyocytes, complete culture medium was exchanged for serum-free medium. Mitotracker Deep Red (40nM; ThermoFisher, UK) was added and incubated for 30 minutes in cell culture conditions. Serum-free medium was then replaced. To image cardiomyocytes as a z stack, beating was stopped by adding 100mM nefidipine (Sigma-Aldrich, Poole, UK) immediately prior to imaging using an Andor Spinning Disk confocal microscope. The Andor Spinning disk system is based on an inverted Olympus IX83 microscope stand and a Yokogawa CSU-X1 spinning disk module. It is equipped with an Oko Labs environmental control chamber to maintain stable conditions for live cell imaging; 37°C, 5% CO_2_ were used throughout. 488nm (BP525/25) and 561nm (LP568) laser lines (and emission filters) operated via an AOTF were used to acquire GFP and mCherry images, respectively. A plan super apochromat 100X 1.4NA oil immersion objective was used throughout, with z steps of 1μm taken. Images were acquired onto an Andor iXon Ultra EMCCD camera (512×512) using an EM gain of 200 with 50ms exposure. Mitochondrial volume was quantified using open-source Mitograph software (Viana *et al.*, 2015; Harwig *et al.*, 2018). This uses 3D reconstructions of labelled organelles to measure the morphology of individual mitochondria as well as characteristics of the mitochondrial network and provides the following outputs: mitochondrial volume, total length and average width. Post-image processing and MitoGraph analysis was performed as per the online protocols (available: http://rafelski.com/susanne/MitoGraph).

### Mitophagy Assay

Primary fetal cardiomyocyte cultures prepared from E14.5-15.5 *mito*-QC fetuses were seeded at a density of 0.25 × 10^6^cells/ml on gelatin-coated glass chambered slides (Ibidi μ-Slide 4 well Glass bottom, Thistle Scientific LTD, Glasgow, UK). After 48 hours, cells were treated with 1μM dexamethasone, 1mM deferiprone (DFP, an iron-chelator, used as a positive control) or vehicle (0.01% ethanol). After 24 hours dexamethasone/DFP/vehicle treated cells were imaged live and z stacks were generated 3, 8 and 45 hours after treatment, using the Andor Spinning Disk confocal live cell imaging system as above. Post-acquisition image analysis was performed blind to the treatment. For each cell, a z stack was acquired and red puncta, indicative of mitophagy, were counted.

### RNA extraction

**Primary fetal cardiomyocytes** seeded in 12 well plates were lysed in 500μl TRIzol (Invitrogen, ThermoFisher, UK) and aspirated into a 1.5ml tube. Chloroform (Sigma-Aldrich, Poole, UK; 100μl) was added and samples vigorously shaken for 30 seconds. Samples were incubated for 2-3 minutes at room temperature then centrifuged 12000 rpm for 15 minutes at 4°C. The aqueous phase was transferred to a new tube containing 250μl isopropanol (Sigma-Aldrich, Poole, UK) and centrifuged as above. The supernatant was discarded and the pellet washed with 500μl 70% ethanol twice. Samples were centrifuged at 10000 rpm for 10 minutes at 4°C, the supernatant removed and pellet air-dried at room temperature for 5-10 minutes before resuspension in 30μl RNAse free water.

**Mouse fetal hearts** were individually homogenised using a stainless steel bead in 500μl RLT buffer (RNeasy, Qiagen, Manchester, UK) and 1% β-mercaptoethanol with a TissueLyser II (Qiagen, Manchester, UK) at maximum speed for 2 minutes. 10μl Proteinase K (20mg/ml; Qiagen, Manchester, UK) and RNAse-free water were added (final volume of 900μl). Samples were incubated at 56°C for 10 minutes, centrifuged at 12000 rpm for 5 minutes at room temperature and the supernatant transferred to a fresh tube containing 400μl 96% ethanol. The mixture was transferred to RNeasy spin tubes (Qiagen mini prep), processed according to the manufacturer’s instructions and eluted in 20μl RNAse free water. This was incubated on the column for at least 5 minutes at room temperature before collection. The first eluate was reapplied to the column and incubated a further minute before the final elution.

**For sheep hearts**, RNA extraction was similar except that samples were minced prior to lysis for 2 sessions each of 3 minutes.

RNA quantity and integrity were determined using a Nanodrop (ThermoFisher, UK) spectrophotometer and gel electrophoresis, respectively.

### Reverse Transcription and quantitative real time PCR

500ng mouse or 300ng sheep RNA was reverse transcribed using QuantiTect Reverse transcription kit (Qiagen, Manchester, UK) and gDNA wipe-out (to remove genomic DNA), according to the manufacturer’s protocol. Included in each sample batch were “no template” and “no reverse transcriptase” controls. Resultant cDNA samples were stored at −20°C. Primers and probes used for qPCR are detailed in Supplementary Table 1. Assays for mouse qPCR were designed using the Roche Universal Probe Library and primers were purchased from Invitrogen (ThermoFisher, UK). A standard curve prepared from pooled cDNA samples was processed with samples on a Lightcycler 480 system (Roche Diagnostics, Burgess Hill, UK). For sheep, Taqman assays (ThermoFisher, UK) were used and performed using a 7900HT Fast Real Time PCR system (ThermoFisher, UK). Internal controls were *Tbp* (for mouse), and *PGK1* and *SDHA* for sheep fetal heart; these did not differ across treatments. To normalise the spread, data were log_10_ transformed.

### Mitochondrial DNA quantification

DNA was extracted from frozen samples using a DNeasy Blood & Tissue kit (Qiagen, Manchester, UK) according to the manufacturer’s protocol. To quantify mitochondrial DNA relative to nuclear DNA, levels of mitochondrial-encoded genes (*Co1, Co2, Nd2*) were measured relative to an intronless nuclear-encoded gene (*Cebpa*) by qPCR (primer sequences in Supplementary Table 1).

### Statistics

Graphpad Prism 8 software was used for statistical analyses. All data are presented as mean ± standard deviation (SD). The number of biological replicates is provided in the figure legends together with the statistical tests used for analysis. All data were subject to Shapiro-Wilk normality testing prior to analysis. Parametric analyses, Student’s t-tests and two-way ANOVA with *post hoc* Sidak’s tests were used as stated in Figure Legends.

## RESULTS

### Dexamethasone increases mitochondrial respiration in primary fetal cardiomyocytes without affecting glycolysis

Consistent with previous findings (Rog-Zielinska *et al.*, 2015), without added fatty acid, basal respiration (oxygen consumption rate; OCR) prior to, and following, addition of 10mM glucose was increased 24 hours following dexamethasone treatment of primary fetal cardiomyocytes (Figure 1A, C). OCR was markedly decreased following addition of oligomycin, an inhibitor of ATP synthase (Figure 1A), suggesting high dependence on mitochondrial respiration for ATP production. As expected, addition of the glycolysis inhibitor, 2-deoxyglucose (2-DG) did not alter the OCR (Figure 1A).

Glycolysis is an important energy source in early fetal cardiomyocytes (Porter *et al.*, 2011). However, our murine primary fetal cardiomyocytes exhibited little dependence on glycolysis. The addition of 2DG did not alter the extra-cellular acidification rate (ECAR; Supplementary Figure 2). In a different approach, using glucose-deprived cells, although the addition of 10mM glucose increased ECAR, this was minimally impacted by 2DG (Figure 1B) and was unaffected by dexamethasone treatment (Figure 1E). This suggests primary fetal cardiomyocytes perform very little glycolysis, relying mainly on mitochondrial oxidation for ATP production.

**Figure 2.**
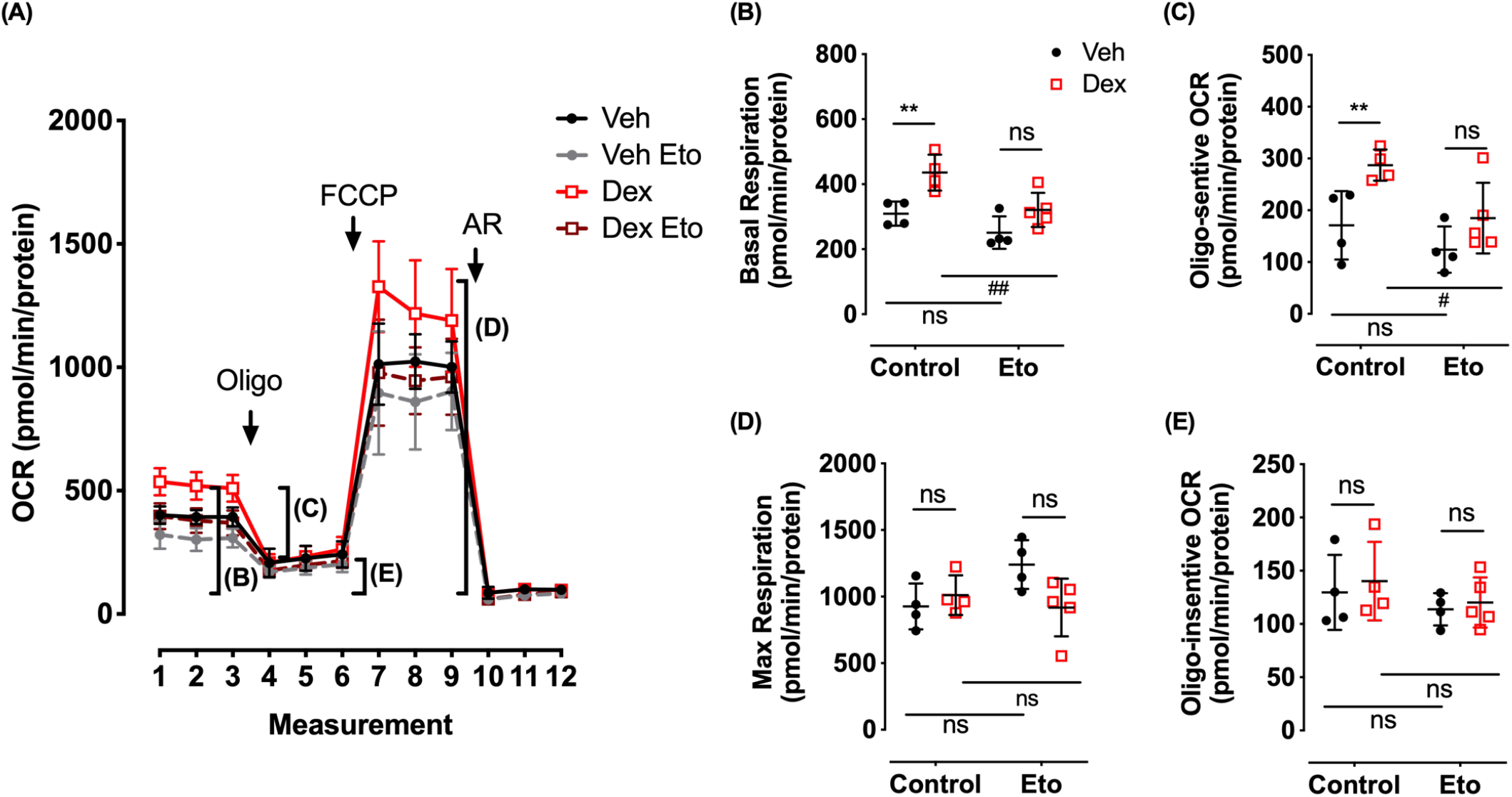
Dexamethasone increases fatty acid oxidation. Primary fetal cardiomyocytes, prepared by digesting pooled E14.5-15.5 C57BL/6J fetal hearts, were cultured for 48 hours in complete DMEM culture medium then treated with dexamethasone (Dex, 1μM, red) or vehicle (Veh, black). After 24 hours, medium was exchanged for Seahorse assay medium supplemented with 5mM glucose, 1mM pyruvate and 0.5mM carnitine. Cells were treated with etomoxir (Eto, 6μM) or vehicle (Control) 15 minutes prior to the addition of BSA-Palmitate (100μM). After a further 15 minutes incubation, cardiomyocyte metabolism was analyzed by extra-cellular flux assay. After 3 basal measurements, oligomycin was added (Oligo, 1.5μM) followed by carbonyl cyanide-4-(trifluoromethoxy)phenylhydrazone (FCCP, 1μM) then antimycin and rotenone (AR, 2μM). Oxygen consumption rate (OCR, **A**) was measured three times over 7.5 minutes following each drug addition. Non-mitochondrial respiration was estimated as the mean OCR remaining after AR addition. (**B**) Basal respiration was estimated as the mean of the 3 basal OCR measurements corrected for non-mitochondrial respiration. (**C**) ATP production was estimated as the maximum change in OCR following the addition of oligomycin. (**D**) Maximum respiration was estimated as the maximum OCR (following FCCP) corrected for non-mitochondrial respiration. (**E**) Leak respiration was estimated as the mean oligomycin-insensitive OCR corrected for non-mitochondrial respiration. Representative data are from an experiment performed on cardiomyocytes pooled from several fetuses across 4-5 wells, mean ± SD, ns=not significant, *p<0.05, ***p<0.001 for comparisons between Veh/Dex, # p<0.05 for comparisons between control and Eto by two-way ANOVA followed by *post hoc* Sidak’s tests. Three samples were excluded due to a technical failure (leakage of AR from the port during basal measurements).

### Dexamethasone increases palmitate oxidation via GR activation

To investigate whether dexamethasone can increase capacity for long chain fatty acid oxidation in fetal cardiomyocytes, OCR was measured in the presence of palmitate. Long chain fatty acids are linked to carnitine and transported into the mitochondrial matrix by carnitine palmitoyltransferase-1 (CPT-1), which is inhibited by etomoxir. Following treatment of primary fetal cardiomyocytes with dexamethasone for 24 hours, in the presence of palmitate there was an increase in basal OCR (Figure 2A, B). Furthermore, in the presence of palmitate, dexamethasone-treated cardiomyocytes showed a larger change in OCR following oligomycin treatment (oligomycin-sensitive OCR; Figure 2C), indicative of increased mitochondrial ATP production. Addition of etomoxir attenuated the dexamethasone-induced increase in basal OCR and mitochondrial ATP production (oligomycin-sensitive OCR) (Figure 2A-C), suggesting that dexamethasone increased palmitate oxidation in fetal cardiomyocytes. Etomoxir itself had no effect on basal respiration or ATP production by fetal cardiomyocytes in the absence of palmitate (Supplementary Figure 1D, E). The dexamethasone-induced increase in fatty acid oxidation was dependent on GR, as pre-treatment of the cardiomyocytes with the GR-antagonist, RU486 blocked the increase in basal respiration and ATP production (Figure 3A-C).

**Figure 3.**
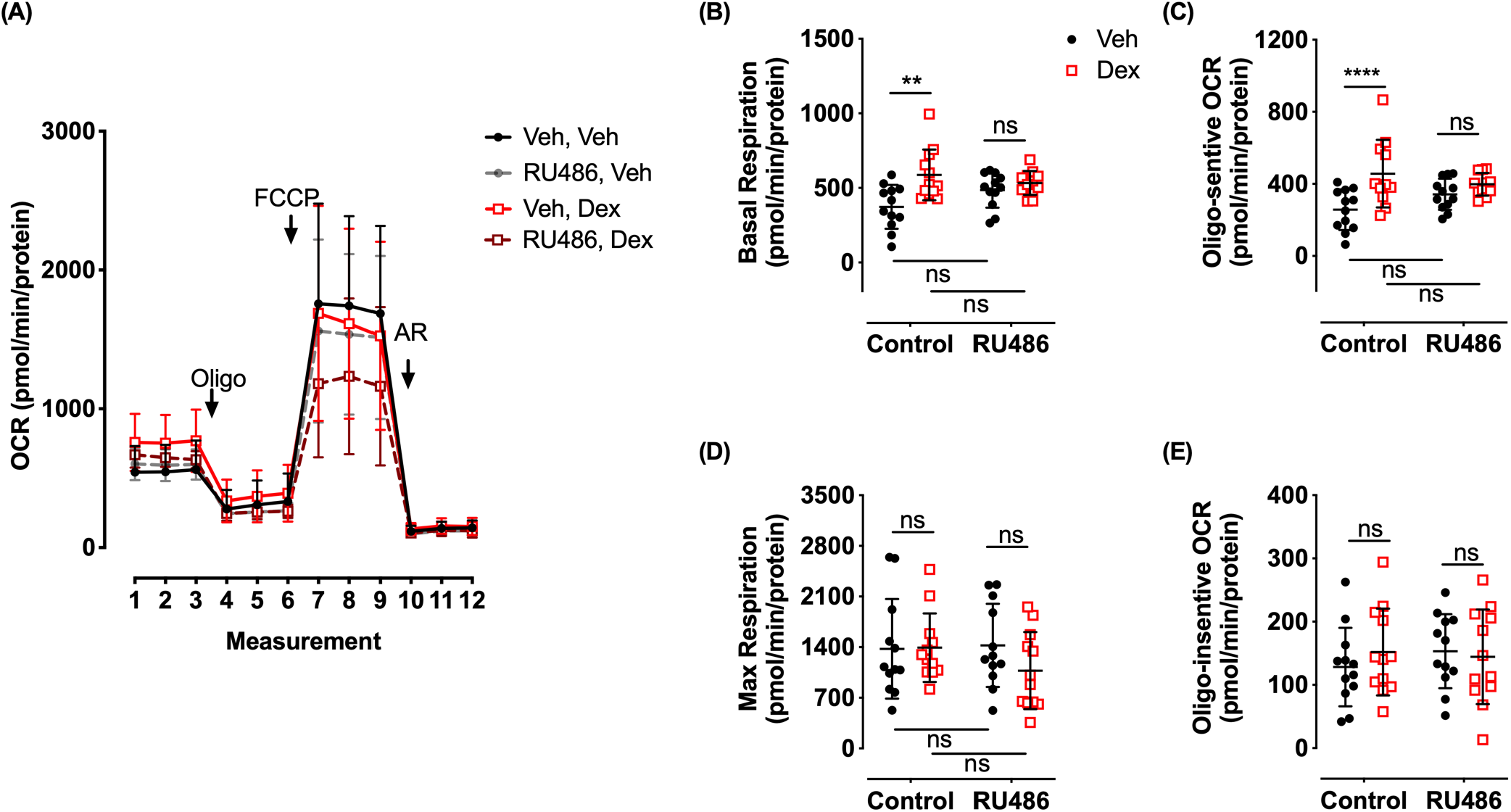
GR mediates the dexamethasone-induced increase in fatty acid oxidation. Primary fetal cardiomyocytes, prepared by digesting pooled E14.5-E15.5 C57BL/6J fetal hearts were cultured for 48 hours before treatment with RU486 (1μM) or vehicle (control) 30 minutes prior to addition of dexamethasone (Dex, 1μM, red) or vehicle (Veh, black). After 24 hours, medium was exchanged for Seahorse assay medium supplemented with 5mM glucose, 1mM pyruvate and 0.5mM carnitine. Cells were treated with etomoxir (Eto, 6μM) or vehicle (Control) 15 minutes prior to addition of BSA-Palmitate (100μM). After 15 minutes incubation with BSA-Palmitate, cardiomyocytes were subjected to extra-cellular flux assay. After 3 basal measurements, oligomycin was added (Oligo, 1.5μM) followed by carbonyl cyanide-4-(trifluoromethoxy)phenylhydrazone (FCCP, 1μM) and antimycin and rotenone (AR, 2μM). Oxygen consumption rate (OCR, **A**) was measured three times over 7.5 minutes following each drug addition. Non-mitochondrial respiration was estimated as the average OCR remaining after AR addition. (**B**) Basal respiration was estimated as the mean of the 3 basal OCR measurements corrected for non-mitochondrial respiration. (**C**) ATP production was estimated as the maximum change in OCR following addition of oligomycin. (**D**) Maximum respiration was estimated as the maximum OCR measurement corrected for non-mitochondrial respiration. (**E**) Leak respiration was estimated as the average oligomycin-insensitive OCR corrected for non-mitochondrial respiration. Data are from cardiomyocytes pooled from tens of fetuses (a total of 12 wells across 3 different pools) and are mean ± SD, ns=not significant, **p<0.01, ****p<0.0001 (two-way ANOVA with *post hoc* Sidak’s tests).

### Dexamethasone upregulates genes involved in long-chain fatty-acid oxidation in fetal cardiomyocytes

The increase in ability to utilise palmitate as a fuel for mitochondrial respiration suggests glucocorticoids increase mitochondrial capacity for long chain fatty acid oxidation. Consistent with the rapid induction of PGC-1α and other master regulators of lipid metabolism in dexamethasone-treated fetal cardiomyocytes (Rog-Zielinska *et al.*, 2015) there was a marked induction of mRNAs encoding enzymes and transporters required for mitochondrial fatty acid oxidation in primary fetal cardiomyocytes 24 hours after treatment with dexamethasone (Figure 4). As well as the master transcriptional regulators, *Ppargc1a* and *Lipin1*, dexamethasone induced expression of *Lcad* and *Mcad* (encoding, respectively, long chain acyl dehydrogenase and medium chain acyl dehydrogenase), *Cd36*, encoding cluster of differentiation-36, also known as fatty acid translocase (a cellular importer of fatty acids), and *Cpt1a* and *Cpt1b*, encoding the alpha and beta subunits, respectively, of CPT-1 (Figure 4). Dexamethasone also increased expression of *Ucp2*, encoding uncoupling protein 2, an inner mitochondrial membrane protein that promotes mitochondrial fatty acid oxidation at the expense of mitochondrial catabolism of pyruvate (Pecqueur *et al.*, 2008). At just 6 hours after addition of dexamethasone, although *Fkbp5*, a well-known glucocorticoid target was strongly up-regulated, levels of *Nr3c1* mRNA, encoding GR, were down-regulated (Supplementary Figure 3), though they recovered by 24 hours (Figure 4). Pre-treatment with RU486 attenuated the dexamethasone-induced increase in *Ppargc1a*, *Lcad*, *Lipin1* and *Cd36* mRNAs (Supplementary Figure 4A-I). At E17.5, hearts of *GR^−/−^* fetal mice had reduced levels of *Mcad* mRNA and a trend for reduced levels of *Ppargc1a* mRNA (Supplementary Figure 4J). The latter is consistent with our previous finding of reduced *Ppargc1a* mRNA in hearts of E17.5 *GR^−/−^* fetal mice (Rog-Zielinska *et al.*, 2013). The failure to reach statistical significance here likely reflects smaller group sizes and over-night matings rather than the time-restricted matings adopted previously.

**Figure 4.**
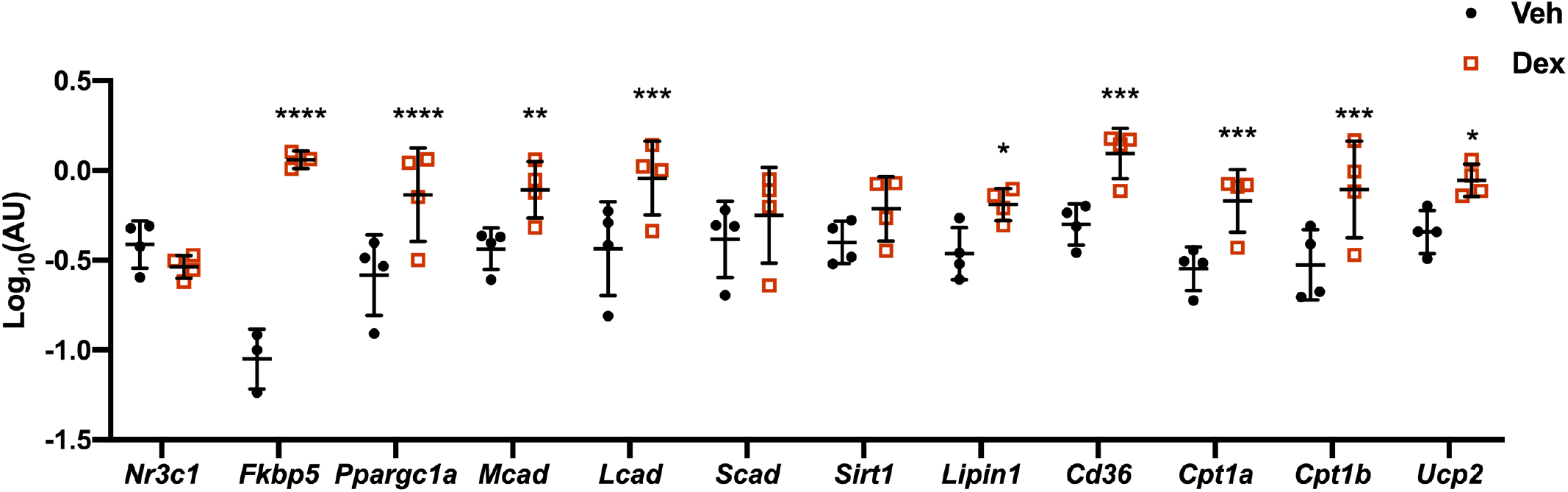
Dexamethasone increases the expression of genes involved in mitochondrial fatty acid oxidation. Primary fetal cardiomyocytes, prepared by digesting pooled E14.5-E15.5 C57BL/6J fetal hearts were cultured for 48 hours before treatment with dexamethasone (Dex, 1μM, red) or vehicle (Veh, black) for 24 hours. Cardiomyocyes were lysed in TRIzol and RNA isolated for analysis by qRT-PCR relative to *Tbp,* used as internal control. Data are from n=4 independent pools of cardiomyocytes, prepared on different days. Data are mean ± SD and were analysed by two-way ANOVA followed by *post hoc* Sidak’s tests; *p<0.05, **p<0.01, ***p<0.001, ****p<0.0001.

### Glucocorticoids do not cause mitochondrial remodelling in fetal cardiomyocytes

Because there is a wave of mitophagy *in vivo* in the mouse fetal heart that coincides with the peak of fetal corticosterone levels (Rog-Zielinska *et al.*, 2013; McWilliams *et al.*, 2016), we hypothesised that glucocorticoids may stimulate the mitophagic replacement of fetal mitochondria by adult mitochondria optimised for fatty acid metabolism (Gong *et al.*, 2015). *mito*-QC transgenic mice utilise a binary fluorescence system in which a ubiquitously expressed tandem mCherry-GFP tag is directed to mitochondria (McWilliams *et al.*, 2016). Under steady-state conditions, the mitochondrial network fluoresces red and green (merged, yellow) in *mito*-QC mice. Upon delivery to lysosomes, the GFP fluorescence, but not that of mCherry, is quenched by the acidic microenvironment. Thus, mitochondria undergoing mitophagic removal appear as punctate mCherry-only foci. To investigate whether dexamethasone induces mitophagy, primary fetal cardiomyocytes from *mito*-QC mice were treated with dexamethasone, vehicle or a mitophagy-inducing agent, deferiprone (DFP) (Allen *et al.*, 2013). As expected, DFP stimulated a robust mitophagy response (as visualised by an increase in the number of mCherry-only positive puncta) over the 45 hour time course (Figure 5A). In contrast, no significant increase in mitophagy could be seen in cardiomyocytes following a similar time course of dexamethasone treatment (Figure 5B). Thus, under these conditions, dexamethasone is not a potent inducer of mitophagy.

**Figure 5.**
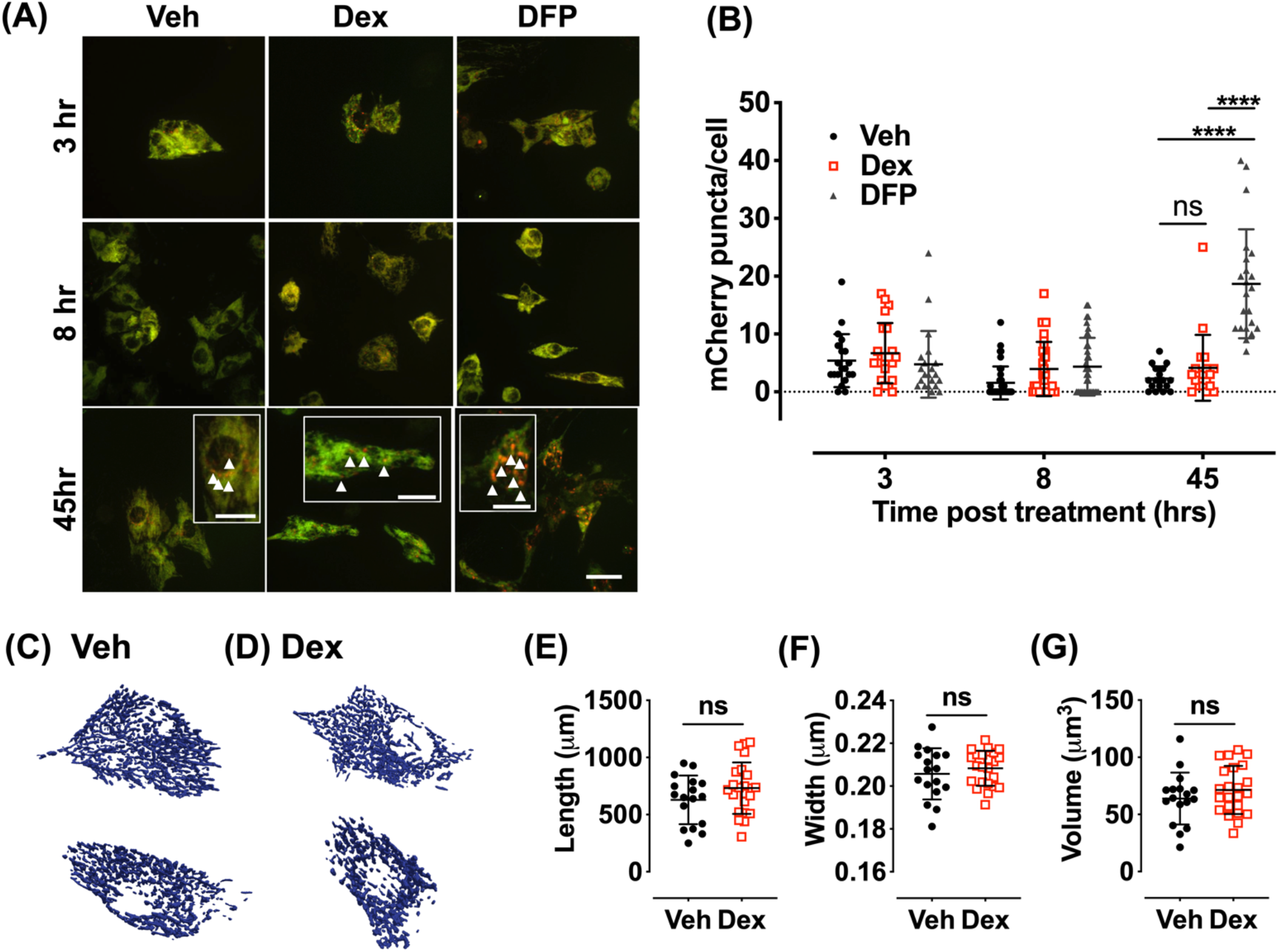
Dexamethasone does not alter mitochondrial morphology or induce mitophagy in primary fetal cardiomyocytes. Primary cardiomyocytes were isolated from E14.5-E15.5 *mito*-QC (**A, B**) or C57Bl/6J (**C-G**) pooled fetal hearts, then cultured for 48 hours. (**A, B**) Cardiomyocytes were then treated with dexamethasone (Dex, 1μM), vehicle (Veh) or a mitophagy inducing agent, deferiprone (DFP, 1mM) and imaged live, 3, 8 and 45 hours later. z stacks of individual cells were obtained using a spinning disk confocal microscope (100X magnification). (**A**) Maximum z projection images are presented: white arrows indicate examples of puncta in magnified panels. Scale bars=10μm and 5μm for main and high magnification panels, respectively. Puncta were counted manually through z stacks (**B**). Data are mean ± SD, n=17-23 individual cells. ****p<0.0001, ns=not significant, two-way ANOVA with *post hoc* Sidak’s tests. (**C-G**) Cardiomyocytes were cultured for 48 hours then treated with dexamethasone (Dex, 1μM) or vehicle (Veh) for 24 hours. Mitochondria were labeled with MitoTracker Deep red CM and cardiomyocyte beating stopped with nefidipine (100mM). z stacks of individual cells were obtained using a spinning disk confocal microscope (100X magnification). Mitochondrial morphology was assessed using MitoGraph software (23, 24) and 3D renderings are shown for vehicle (**C**) and dexamethasone (**D**) treated cardiomyocytes. Parameters included (**E**) length, (**F**) width and (**G**) total volume. Data are mean ± SD, n=17-40 individual cells. ns=not significant (t-tests).

Glucocorticoids have been associated with changes in mitochondrial number and function (Weber *et al.*, 2002; Du *et al.*, 2009; Lapp *et al.*, 2018). Accordingly, we next investigated whether the dexamethasone-induced increase in fatty acid oxidation was associated with an increase in mitochondrial volume and/or number. Following dexamethasone treatment of primary fetal cardiomyocytes for 24 hours, mitochondria were labelled with Mitotracker Deep Red CM and the mitochondrial network imaged (Figure 5C, D). There were no differences in mitochondrial volume, length or width between dexamethasone and vehicle treated cardiomyocytes (Figure 5E-G). Similarly, measurements of the GFP-fluorescent mitochondrial network in fetal cardiomyocytes from *mito*-QC mice showed no differences in mitochondrial morphology as a result of dexamethasone treatment (Supplementary Figure 5A-C). Furthermore, mitochondrial DNA content, an indirect measurement of mitochondrial number, did not differ between hearts of E17.5 *GR^−/−^* mice and their control *GR^+/+^* littermates (Supplementary Figure D-F). Thus, the glucocorticoid-mediated increase in fatty acid oxidation capacity most likely occurs independently of any change in mitochondrial number or morphology.

### *In vivo*, dexamethasone-induced changes in fatty acid oxidation genes in mouse hearts are developmental stage-dependent

To investigate whether glucocorticoid administration *in vivo* can similarly induce cardiac fatty acid oxidation capacity, dexamethasone or vehicle was administered to pregnant dams at E13.5 or E16.5 or to neonatal mice at postnatal day (P)1. Hearts were examined 24 hours after injection. E14.5 fetal hearts appeared glucocorticoid-resistant: dexamethasone had no significant effect on any of the mRNAs examined, including the glucocorticoid target, *Fkbp5*, and *Nr3c1* encoding GR itself (Figure 6A). However, at E17.5, dexamethasone downregulated cardiac *Nr3c1* mRNA levels (Figure 6B). At the same time, levels of *Ppargc1a* and mRNA encoding enzymes and transporters for fatty acid oxidation (*Mcad, Lcad, Lipin1, Cd36, Cpt1a, CPT1b*) were strongly downregulated, despite a trend for a modest increase in *Fkbp5* mRNA (Figure 6B). In complete contrast, at P2, *Fkbp5* was strongly induced by dexamethasone, as was *Ppargc1a* and the fatty acid oxidation genes: *Mcad, Lcad, Cd36, Cpt1b* (Figure 6C). Moreover, levels of *Nr3c1* mRNA were unchanged (Figure 6C). Thus, the effect of glucocorticoids upon cardiac capacity for fatty acid oxidation reflect the effect upon GR expression itself and its key target gene, *Ppargc1a.* To explore whether this regulation extends to a more translationally relevant model, we measured cardiac mRNA encoding GR and PGC-1α in a sheep model of preterm birth following antenatal corticosteroid administration that mimics current clinical practice. Celestone is a mix of betamethasone phosphate and betamethasone acetate that is widely used as an antenatal corticosteroid in the USA, Europe (though not the UK), Australia and New Zealand. Preterm lambs delivered at 127 days (term being ~147 days) 48 hours after initiating a course of Celestone (2 doses, administered 24 hours apart) showed reduced expression of *NR3C1* mRNA in both left and right ventricles of the heart (Figure 7A, B). Administration of a single dose of betamethasone acetate (equivalent to just the betamethasone acetate component of Celestone) 24 hours before delivery also reduced *NR3C1* mRNA levels, though this did not achieve significance in the right ventricle (Figure 7A, B). Levels of *PPARGC1* mRNA were reduced in the right ventricle following Celestone administration, with a more modest effect (p>0.05) in the left ventricle. Betamethasone acetate alone caused a similar though non-significant reduction in *PPARGC1* mRNA levels in both ventricles (Figure 7C, D). This suggests that antenatal corticosteroid administration may interfere with the normal maturation of the mid to late gestation heart by down-regulating GR.

**Figure 6.**
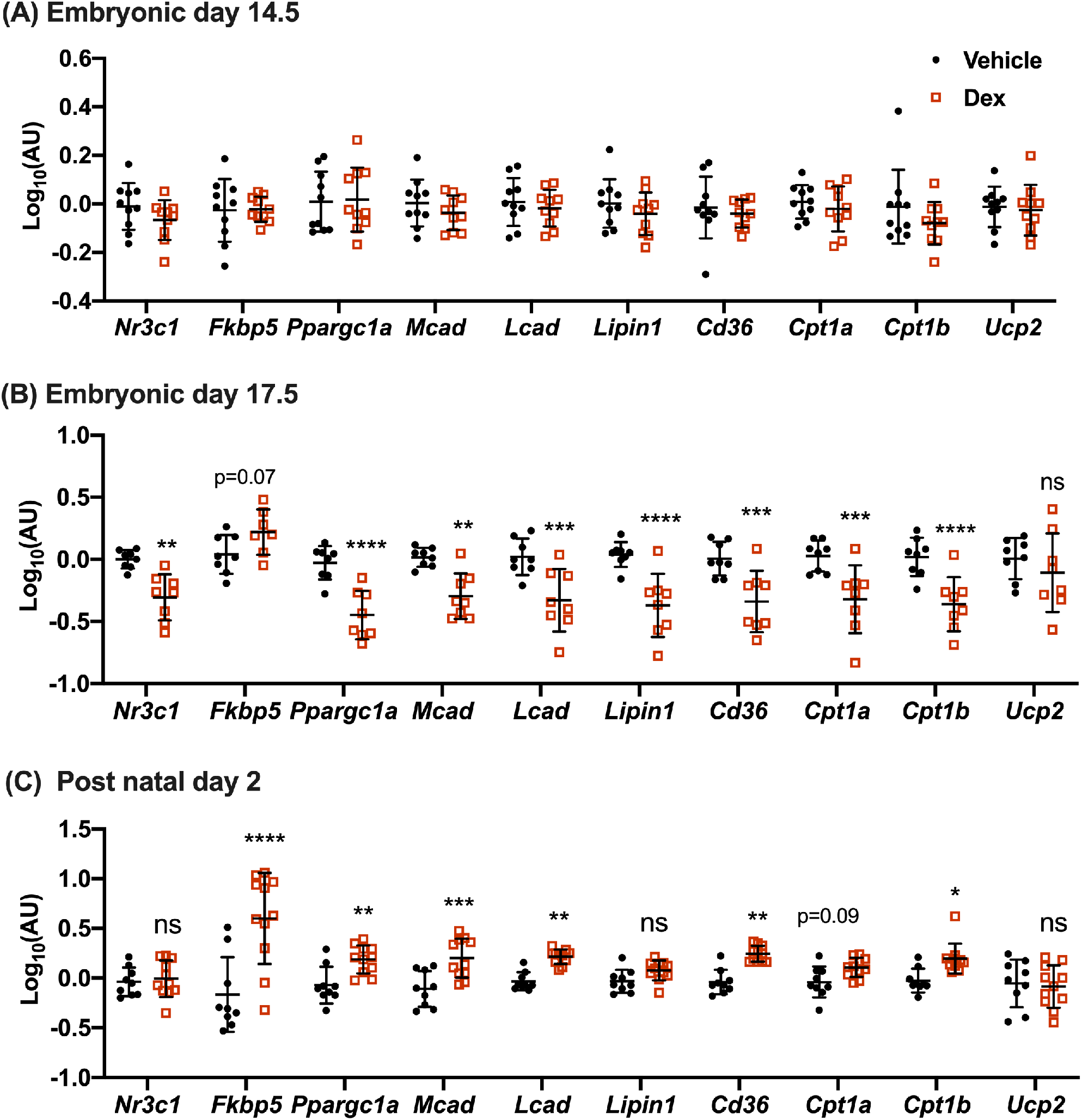
Dexamethasone regulates cardiac mitochondrial fatty acid oxidation in perinatal mice in vivo. Pregnant C57Bl/6J dams were injected (i.p.) with 0.5mg/kg dexamethasone or vehicle at E13.5 (**A**) or E16.5 (**B**). C57Bl/6J neonates (**C**) were injected on P1 with dexamethasone (0.5mg/kg) or vehicle. After 24 hours hearts were excised and analysed by qRT-PCR for genes involved in fatty acid oxidation. Data are from n=8-10 individual animals from at least 5 litters/group. Data are mean ± SD and were analysed by two-way ANOVA followed by *post hoc* Sidak’s tests; *p<0.05, **p<0.01, ***p<0.001, ****p<0.0001.

**Figure 7.**
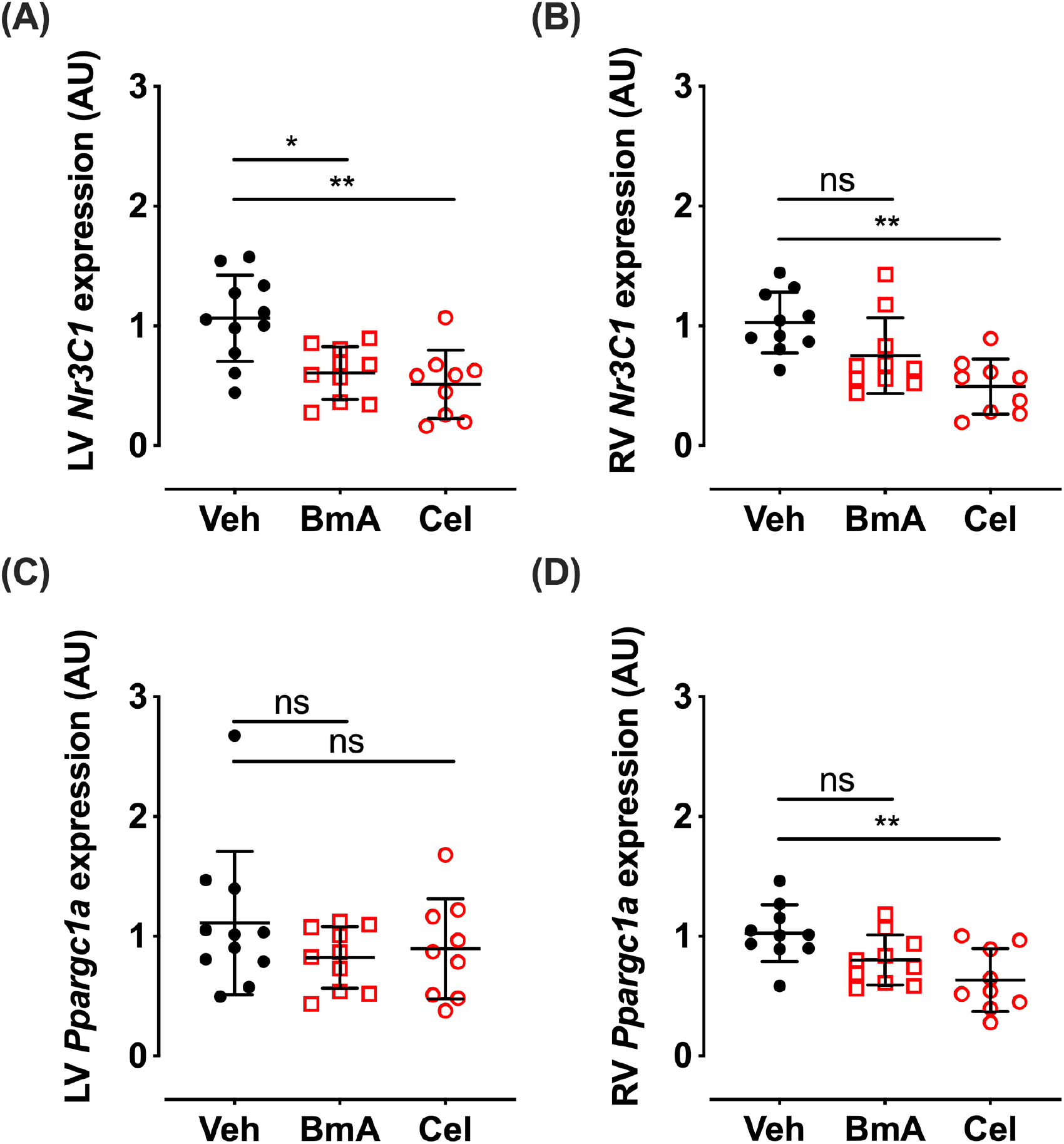
Cardiac GR and PGC-1α expression are reduced following treatment with antenatal corticosteroids in a sheep model of pre-term birth. Lambs were delivered preterm at 122±1 days (term being ~147 days), 48 hours after initiating a course of celestone (Cel: 2 doses, 24 hours apart) or 24 hours after a single dose of betamethasone acetate (BmA). (**A, B**) mRNA encoding GR (*NR3C1*) and (**C, D**) PGC-1α (*PPARGC1A*) were measured by qPCR in the left ventricle (LV) and right ventricle (RV). Data are mean ± SD, ns=not significant, *p<0.05, **p<0.01, analysed by one way ANOVA with *post hoc* Tukey’s tests. For RV: Veh n=10, BmA n=10, Cel n=9. For LV: Veh n=11, BmA n=10, Cel n=9.

## DISCUSSION

Here, we find that mouse fetal cardiomyocytes use mainly mitochondrial metabolism to generate ATP when glucose is provided as substrate, with little reliance on glycolysis. This supports the view that metabolism has switched from anaerobic glycolysis to aerobic mitochondrial respiration by the end of the embryonic period at E14.5 in mice (reviewed (Porter *et al.*, 2011)). Although glucocorticoids increase basal mitochondrial respiration (confirming previous findings (Rog-Zielinska *et al.*, 2015)), they have no effect on glycolysis, ruling out a glucocorticoid-promoted switch from glycolysis to oxidative metabolism. Our data do not support a glucocorticoid-mediated increase in mitochondrial number or change in morphology to account for the increase in basal respiration. Instead, our data suggest glucocorticoid action in the fetal heart promotes mitochondrial ATP generating capacity, in line with their maturational effects.

As well as increasing basal mitochondrial respiration with carbohydrate substrates, glucocorticoid treatment of fetal cardiomyocytes *in vitro* increases fatty acid oxidation. It has been suggested that mitophagy is required to replace the mitochondrial network in perinatal cardiomyocytes with mitochondria optimised for fatty acid oxidation (Gong *et al.*, 2015). However, our data suggest that an increase in fatty acid oxidation in perinatal cardiomyocytes can occur without substantial mitophagy. Moreover, they are consistent with the notion that mitochondrial remodelling occurs in cardiomyocytes prior to our cell isolations at ~E15. Mitochondrial phenotype in the embryonic heart changes considerably between E9.5 and E13.5, compatible with mitochondrial remodelling by mitophagy being required for the switch in cardiomyocyte reliance from anaerobic glycolysis to aerobic respiration. By E13.5, the network is interconnected and spans the cell, more closely resembling that in late fetal cardiomyocytes (Porter *et al.*, 2011). Our conclusions differ from a recent report that suggested that dexamethasone promotes mitophagy in mouse embryonic-stem cell-derived cardiomyocytes through Parkin (Zhou *et al.*, 2020). However, in those experiments, detection of lysosomes (using lysotracker) was only possible in dexamethasone-treated cells (Zhou *et al.*, 2020) making interpretation of the mitophagy findings difficult. Nevertheless, dexamethasone did not affect mitochondrial morphology in mouse embryonic-stem cell-derived cardiomyocytes (Zhou *et al.*, 2020), consistent with our findings here. Our data reported here clearly show that although glucocorticoid treatment in primary mouse fetal cardiomyocytes increases fatty acid oxidation, it does not induce widespread mitophagy.

Fatty acid oxidation dramatically increases around the end of the first postnatal week in mouse heart, and we found a marked induction of the pathway following glucocorticoid administration *in vivo* in neonatal mice (Lopaschuk & Jaswal, 2010). However, in preparation for postnatal life, fatty acid oxidation has already begun to occur in the late gestation fetal heart (Lopaschuk & Jaswal, 2010; Porter *et al.*, 2011). In sheep, there is an increase in cardiac expression of genes related to fatty acid oxidation between late gestation and term that continues after birth (Richards *et al.*, 2015). *In silico* transcription factor analysis suggests this is, at least in part, GR-mediated (Richards *et al.*, 2015). The lower level of *Mcad* mRNA in hearts of GR knockout fetuses at E17.5, a stage when endogenous glucocorticoid levels have increased, supports a role for GR in the late gestation increase in cardiac expression of genes required for fatty acid oxidation. Fatty acid oxidation genes themselves are not primary targets of GR in mouse fetal cardiomyocytes (Rog-Zielinska *et al.*, 2015) and are likely indirectly regulated by master regulators of fatty acid oxidation including CEBPβ, PPARα and PGC-1α, which are primary GR targets (Rog-Zielinska *et al.*, 2015). Thus, *endogenous* glucocorticoid action, via GR, may contribute to the normal rise in fatty acid oxidation capability as the fetus approaches term. This is likely to be important to meet the increase in cardiac energy demand after birth, consistent with the ergogenic effects of glucocorticoids (Addison, 1855; Morrison-Nozik *et al.*, 2015) and the vital role of the late gestation increase in glucocorticoids to prepare for life after birth.

Crucially, our data illustrate how *exogenous* glucocorticoid may interfere with the normal maturation of energy metabolism in the fetal heart. Although dexamethasone increases expression of fatty acid oxidation genes in neonatal mice, 24 hours after administration of dexamethasone at E16.5 (the peak of endogenous fetal glucocorticoid levels) the expression of genes related to fatty acid oxidation actually *decreases* in the fetal heart. The association with reduced *Nr3c1* mRNA, encoding GR itself, suggests the decrease in the mitochondrial fatty acid oxidation pathway reflects down-regulation of glucocorticoid signalling *per se* in the fetal heart following dexamethasone treatment. We recently reported similar down-regulation of GR expression in fetal heart as well as reduced endogenous fetal corticosterone levels following dexamethasone treatment (via drinking water) between E12.5-E15.5 (Agnew *et al.*, 2019). This was associated with a transient alteration in fetal diastolic heart function (Agnew *et al.*, 2019). Whilst dexamethasone also down-regulates *Nr3c1* mRNA in fetal cardiomyocytes *in vitro*, this is transient with recovery of *Nr3c1* mRNA expression by 24 hours. In the neonatal heart, if *Nr3c1* mRNA is transiently downregulated by dexamethasone, it has recovered within 24 hours. Dynamic and differential auto-regulation of GR, previously described for adult tissues (Kalinyak *et al.*, 1987; Spencer *et al.*, 1991; Freeman *et al.*, 2004), may contribute to the complex and context dependent effects of perinatal glucocorticoid administration. Previous studies have examined the effect of antenatal glucocorticoids in rodents and reported contradictory findings (reviewed, (Rog-Zielinska *et al.*, 2014)). Our data illustrate that exogenous glucocorticoids can potentially interfere with normal heart maturation. Indeed, maturation of the rat heart is delayed after prenatal treatment with dexamethasone (Torres *et al.*, 1997). Timing may be critical. Although antenatal dexamethasone increased ATP content in the neonatal rat heart, the same treatment did not increase cardiac ATP content prior to birth, despite an increase in creatine kinase expression (Mizuno *et al.*, 2010). GR expression was not examined in that study. Investigations in sheep have also highlighted adverse effects of antenatal glucocorticoid exposure on cardiac energy metabolism. Maternal hypercortisolaemia reduces fetal cardiac mitochondrial number and oxidative metabolism at term, associated with fetal ECG abnormalities, an inability to maintain fetal aortic pressure and heart rate during labour and a dramatic increase in perinatal death (Antolic *et al.*, 2018). Again, whether GR expression was affected was not reported.

Differential mRNA stability may explain some of the complex effects of glucocorticoids on downstream genes. PGC-1α is essential for efficient and maximal fatty acid oxidation and ATP production in cardiomyocytes (Arany *et al.*, 2005; Lehman *et al.*, 2008). *Ppargc1a* mRNA has a short half-life, being less than 30 minutes in rat skeletal muscle extracts. This is further decreased with chronic muscle stimulation (Lai *et al.*, 2010). Consistent with a short half-life, we have previously shown that blocking new protein synthesis with cycloheximide increases *Ppargc1a* mRNA levels in fetal cardiomyocytes (Rog-Zielinska *et al.*, 2015), suggesting it is actively degraded. Glucocorticoid treatment rapidly increases levels of *Ppargc1a* mRNA in fetal heart *in vivo* and in fetal cardiomyocytes *in vitro* (Rog-Zielinska *et al.*, 2015). Moreover, dexamethasone "super-induces" *Ppargc1a* in the presence of cycloheximide (Rog-Zielinska *et al.*, 2015), again consistent with a rapid turnover of *Ppargc1a* mRNA. A very short half-life and a need for activated GR to continually enhance transcription of *Ppargc1a* mRNA may explain the close association between *Nr3c1* and *Ppargc1a* mRNA in both fetal heart and mouse primary cardiomyocytes that we saw here. By contrast, in all of our experiments, with the exception of fetal heart at E14.5 (which appeared glucocorticoid resistant), *Fkbp5* mRNA was upregulated by dexamethasone. Even in the E17.5 heart, there was a strong trend for increased *Fkbp5* mRNA following dexamethasone, despite downregulation of GR and the fatty acid oxidation pathway at this time. Just 6 hours after addition of dexamethasone to fetal cardiomyocytes *in vitro*, *Fkbp5* mRNA was already elevated, despite downregulation of GR. Thus, *Fkbp5* mRNA appears a stable readout of early GR activation whereas *Ppargc1a* mRNA correlates with *Nr3c1* mRNA at any particular time. Plausibly, antentatal corticosteroids may disrupt the normal maturation of energy metabolism in the human fetal heart, in part by down-regulating PGC-1α. In our sheep model that closely mirrors clinical practice, both *NR3C1* and *PPARGC1A* were downregulated in fetal heart by antenatal corticosteroids. Whether this is transient, and both later recover to control levels merits further investigation. Nevertheless, this suggests that clinical administration of antenatal corticosteroids in mid- and late gestation may interfere with normal heart maturation.

The E13.5 fetal heart appears resistant to dexamethasone. The reason for this is unclear, but it has implications for clinical practice. In humans, the reduction in fetal heart rate variability (a clinical marker of fetal hypoxia/poor outcomes) 2 to 3 days after maternal administration of glucocorticoids is greater in fetuses >30 weeks gestation than those <30 weeks (Mulder *et al.*, 2009), consistent with gestation-stage dependent effects of antenatal glucocorticoids. The greater sensitivity to the haemodynamic effects of glucocorticoids coincides with an increase in fetal cortisol synthesis, from ~30 weeks gestation (Hillman *et al.*, 2012). In mice, adrenal steroidogenesis initiates at E14.5 (Michelsohn & Anderson, 1992). This raises the possibility that the fetus is glucocorticoid resistant prior to the gestational increase in fetal glucocorticoid levels. Whether this is the case or not merits future investigation, given wide-spread expression of GR in the mouse fetus prior to E14.5. It is interesting to note that although dexamethasone increases calcium handling and contraction force in human embryonic stem cell-derived cardiomyocytes (Kosmidis *et al.*, 2015), human induced pluripotent stem cell-derived cardiomyocytes (roughly corresponding to first trimester human fetal cardiomyocytes (van den Berg *et al.*, 2015)) do not respond to dexamethasone alone, possibly because they lack a competence factor (Birket *et al.*, 2015). The acquisition of competence to respond to glucocorticoids as well as the auto-regulation of GR itself may therefore be developmentally regulated and may differ between cell types. Understanding how this contributes to the maturational effects of glucocorticoids upon fetal organs and tissues will be vital to optimise antenatal corticosteroid therapy in the future, to limit possible harm and maximise benefit.

## ABBREVIATIONS

GR: glucocorticoid receptor
E: embryonic day
P: postnatal day
PBS: phosphate buffered saline
SRB: sulforhodamine B
2DG: 2 deoxyglucose
AR: antimycin and rotenone
OCR: oxygen consumption rate
ECAR: extracellular acidification rate
FCCP: carbonyl cyanide-4-(trifluoromethoxy)phenylhydrazone
BSA: bovine serum albumin
GFP: green fluorescent protein
DFP: deferiprone
CPT-1: carnitine palmitoyltransferase-1
PGC-1α – PPARγ: coactivator-1α
SD: standard deviation

## ACKNOWLEDGEMENTS

We are grateful to staff at The University of Edinburgh Central Bioresearch Services for assistance with animal care, particularly Hollie McGrath and Sandra Spratt. We thank colleagues in the Centre for Cardiovascular Science, especially Megan Holmes, Martin Denvir and Gillian Gray for helpful discussions. We are grateful to Merck & Co. for the gift of betamethasone acetate.

## DATA AVAILABILITY

The data that support the findings of this study are available from the first and/or corresponding author upon reasonable request.

## AUTHOR CONTRIBUTIONS

Conceptualisation – JRI, EAR-Z, KEC; data curation – JRI, KEC, SJS; data analysis – JRI, LH, RNC, NMM; Funding – KEC, IGG, SJS, MWK, NMM, CW; Investigation – JRI, RNC, J-FZ, CB, HU, EAR-Z, EP, LH, CN, SJS, MWK; Methodology – JRI, RNC, J-FZ, CB, EAR-Z, EJA, MWK; Project administration – JRI, KEC; Resources – IGG, MWK, NMM, KEC; Supervision – JRI, KEC; Validation – JRI, KEC; Writing – original draft – JRI, MWK, KEC; Writing – review and editing – JRI, RNC, CB, EAR-Z, EJA, MWK, SJS, IGG, KEC. All authors approved the final version of the manuscript, agree to be accountable for all aspects of the work in ensuring that questions related to the accuracy or integrity of any part of the work are appropriately investigated and resolved and qualify for authorship. All those who qualify for authorship are listed.

