## Supplementary information for "GLUCOCORTICOIDS REGULATE MITOCHONDRIAL FATTY ACID OXIDATION IN FETAL CARDIOMYOCYTES"

***SUPPLEMENTARY TABLE 1*. Sequences of primers and probes used in real-time PCR**

Forward and reverse primers and Roche universal probe library (Invitrogen, ThermoFisher, UK) probe number used in real-time PCR assays (mouse, unless specified otherwise) to measure specific cDNA/mitochondrial DNA levels.

| Assay | Comment | accession number | forward primer 5'-3' | reverse primer 5'-3' | UPL probe |
|---|---|---|---|---|---|
| *Rn18s* | 18S RNA | NR_003278.1 | CTCAACACGGGAAACCTCAC | CGCTCCACCAACTAAGAACG | 77 |
| *Tbp* | TATA box binding protein | NM_013684 | GGGAGAATCATGGACCAGAA | GATGGGAATTCCAGGAGTCA | 97 |
| *Nr3c1* | Glucocorticoid receptor (mouse) | NM_008173.4 | TGACGTGTGGAAGCTGTAAAGT | CATTTCTTCCAGCACAAAGGT | 56 |
| *Fkbp5* | FK509 binding protein | NM_010220.4 | AAACGAAGGAGCAACGGTAA | TCAAATGTCCTTCCACCACA | 97 |
| *Ppargc1a* | PGC-1$\alpha$ (mouse) | NM_008904.2 | GAAAGGGCCAAACAGAGAGA | GTAAATCACACGGCGCTCTT | 29 |
| *Acadm (Mcad)* | Medium chain specific acyl-CoA dehydrogenase | NM_007382.5 | GGAAAGCTGCTAGTGGAGCA | TGCAATCGAGGCATAGTAAGTG | 64 |
| *Acadl (Lcad)* | Long chain specific acyl-CoA dehydrogenase | NM_007381.4 | AGTGTATCGGTGCCATAGCC | TGGCGTTCGTTCTTACTCCT | 11 |
| *Lpin1 (Lipin1)* | Lipin1 | NM_172950.3 | GACTGGGAAAGGCCACAATA | CGTGCTCTTCATCACTGGAG | 49 |
| *Cd36* | CD36 | NM_001159555 | TTGAAAAGTCTCGGACATTGAG | TCAGATCCGAACACAGCGTA | 6 |
| *Cpt1a* | Carnitine pamitoyltransferase-1$\alpha$ | NM_013495.2 | CTCAGTGGGAGCGACTCTTC | TGTCCTTGACGTGTTGGATG | 94 |
| *Cpt1b* | Carnitine pamitoyltransferase-1$\beta$ | NM_009948.2 | GTACCGCCTAGCCATGACA | GGCTCCAGGGTTCAGAAAGT | 29 |
| *Ucp2* | Uncoupling protein 2 | NM_011671.5 | ACAGCCTTCTGCACTCCTG | GGCTGGGAGACGAAACACT | 2 |
| *Mt-CO1* | Mitochondrial encoded cytochrome oxidase-1 (mitochondrial genomic DNA) | N/A | CAGACCGCAACCTAAACACA | TTCTGGGTGCCCAAAGAAT | 25 |
| *Mt-CO3* | Mitochondrial encoded cytochrome oxidase-2 (mitochondrial genomic DNA) | N/A | TAGCCTCGTACCAACACATGA | AGTGGTGAAATTCCTGTTGGA | 66 |
| *Mt-ND2* | Mitochondrial encoded NADH dehydrogenase-2 (mitochondrial genomic DNA) | N/A | CAAATTTACCCGCTACTCAACTC | GCTATAATTTTTCGTATTTGTGTTTGG | 101 |
| *Cebpa* | Nuclear encoded C/EBP$\alpha$ (intronless gene) | TaqMan ABI Mm_00514283_s1 | | | |
| *NR3C1* | Glucocorticoid receptor (sheep) | TaqMan ABI Oa_04657790_m1 | | | |
| *PPARGC1A* | PGC-1$\alpha$ (sheep) | TaqMan ABI Oa_01208835_m1 | | | |
| *PGK1* | Phosphoglycerate kinase-1 (sheep) | TaqMan ABI Oa_04657427_gH | | | |
| *SDHA* | Succinate dehydrogenase subunit A | TaqMan ABI Oa_04307499_m1 | | | |

**Ivy et al, Supplementary Figure 1**

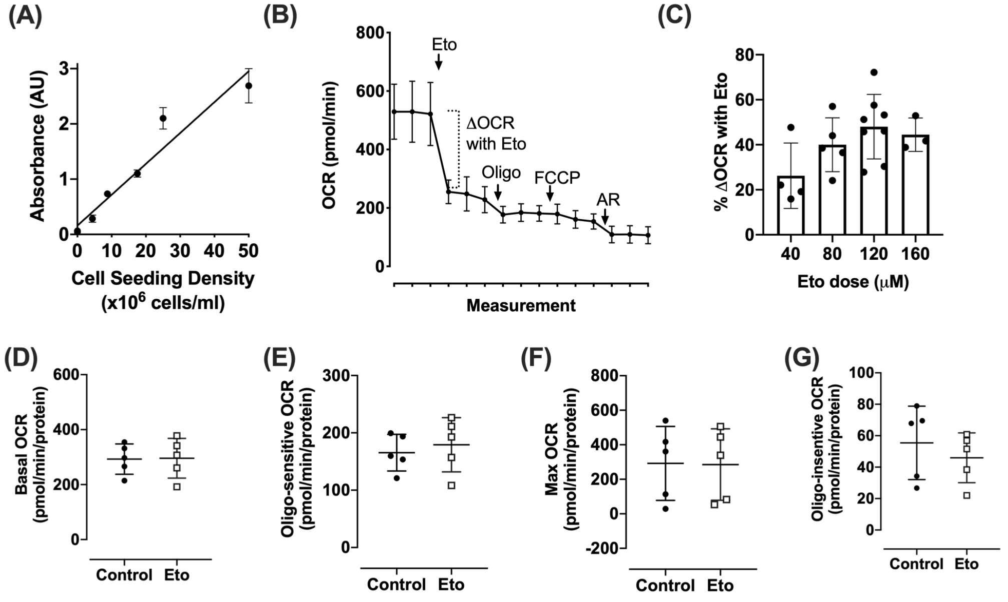


***SUPPLEMENTARY FIGURE 1.*** **Optimisation experiments for Extra-cellular Flux Assays in Primary Fetal Cardiomyocytes**

Primary fetal cardiomyocytes were prepared by digesting pooled E14.5-15.5 C57BL/6J fetal hearts then cultured for 48h in complete DMEM culture medium. (A) Sulpho-rhodamine assays, used to estimate protein levels for normalization of extra-cellular flux assays, were linear over a range of primary fetal cardiomyocyte seeding densities: data from a single pool of cardiomyocytes cultured in triplicate wells, line fitted by linear regression with R-square value of 0.93. (B) Cardiomyocytes were treated with dexamethasone (Dex, 1μM) or vehicle (Veh, 0.01% ethanol) for 24h, then medium was exchanged for seahorse assay medium supplemented with 5mM glucose, 1mM pyruvate and 0.5mM carnitine. Cardiomyocytes were treated with BSA-Palmitate (100μM) 30min prior to extra-cellular flux analysis. After 3 basal measurements, Etomoxir was added (Eto, 120μM), followed by oligomycin (Oligo, 1.5μM), carbonyl cyanide-4-(trifluoromethoxy)phenylhydrazone (FCCP, 1μM) and antimycin and rotenone (AR, 2μM). Following treatment with 120μM Etomoxir, cells were unable to respond to the uncoupling agent, FCCP. There was a very small response to AR. Following a literature review, we concluded that high doses of Etomoxir may inhibit Complex I and III at high doses and thus a lower dose (6μM) was selected for future experiments. (C) Cardiomyocytes were treated with BSA-Palmitate (100μM) in seahorse assay medium 30min prior to extra-cellular flux analysis. Oxygen consumption rate (OCR), measured following sequential addition of increasing doses of etomoxir (Eto), indicated that 120μM gave the maximum change in oxygen consumption rate (OCR), with a plateau thereafter; n=3-5 technical replicates over 2 separate experiments. (D-G) The effect of Etomoxir (6μM) in the absence of the fatty, acid palmitate was investigated. Cardiomyocytes were treated with dexamethasone (Dex, 1μM) or vehicle (Veh, 0.01% ethanol) for 24h, then cell culture medium was exchanged for seahorse assay medium supplemented with 5mM glucose and 0.5mM carnitine. Cells were treated with etomoxir (Eto, 6μM) or vehicle (Control, medium) 15min prior to the addition of BSA alone (17μM). After 15min incubation, metabolism was analyzed by extra-cellular flux assays. After 3 basal measurements, oligomycin was added (Oligo, 1.5μM) followed by carbonyl cyanide-4-(trifluoromethoxy)phenylhydrazone (FCCP, 1μM) and antimycin and rotenone (AR, 2μM). Oxygen consumption rate (OCR) was measured in triplicate for 2.5min following each drug addition. Non-mitochondrial respiration was estimated as the average OCR remaining after treatment with AR. (D) Basal respiration was estimated as the mean of the 3 basal OCR measurements corrected for non-mitochondrial respiration. (E) ATP production was estimated as the maximum change in OCR following the addition of oligomycin (oligomycin-sensitive OCR). (F) Maximum respiration was estimated as the maximum OCR measurement induced by FCCP corrected for non-mitochondrial respiration. (G) Leak respiration was estimated as the mean oligomycin-insensitive OCR corrected for non-mitochondrial respiration. Data are mean ± SD, ns= not significant, data analysed by two-way ANOVA, n=5 technical replicates.

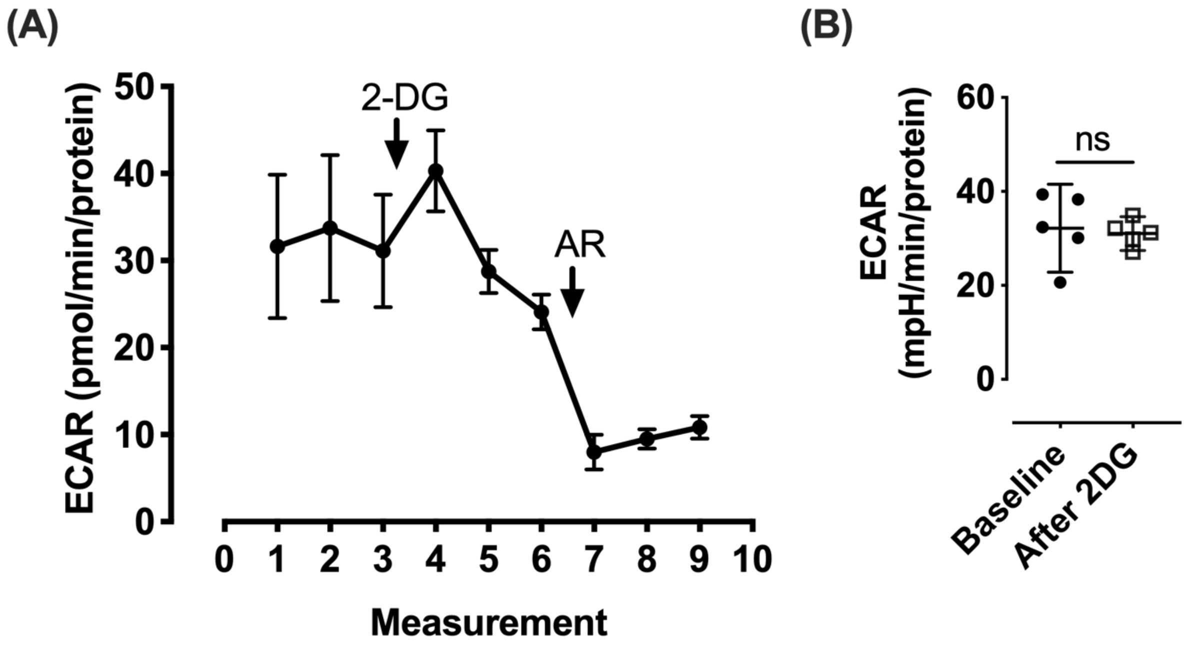


***SUPPLEMENTARY FIGURE 2.*** **Primary fetal cardiomyocytes exhibit negligible glycolysis**

(A) Primary fetal cardiomyocytes, prepared by digesting pooled E14.5-15.5 C57BL/6J fetal hearts were cultured for 72h in complete DMEM culture medium then exchanged for seahorse base medium supplemented with 10mM glucose, 1mM pyruvate. After 3 baseline measurements 2-deoxyglucose (2DG, 100mM) was added followed by antimycin and rotenone (AR, 2μM). 3 measurements at 2.5min intervals were recorded after each addition. (B) The maximum difference between extra-cellular acidification rate (ECAR) at baseline (Measurement 2) and after addition of the glycolysis inhibitor, 2-DG (Measurement 6) was used as a measure of glycolysis. Data are mean ± SD, Data analysed by Student's t-tests where ns= not significant, n=5 technical replicates.

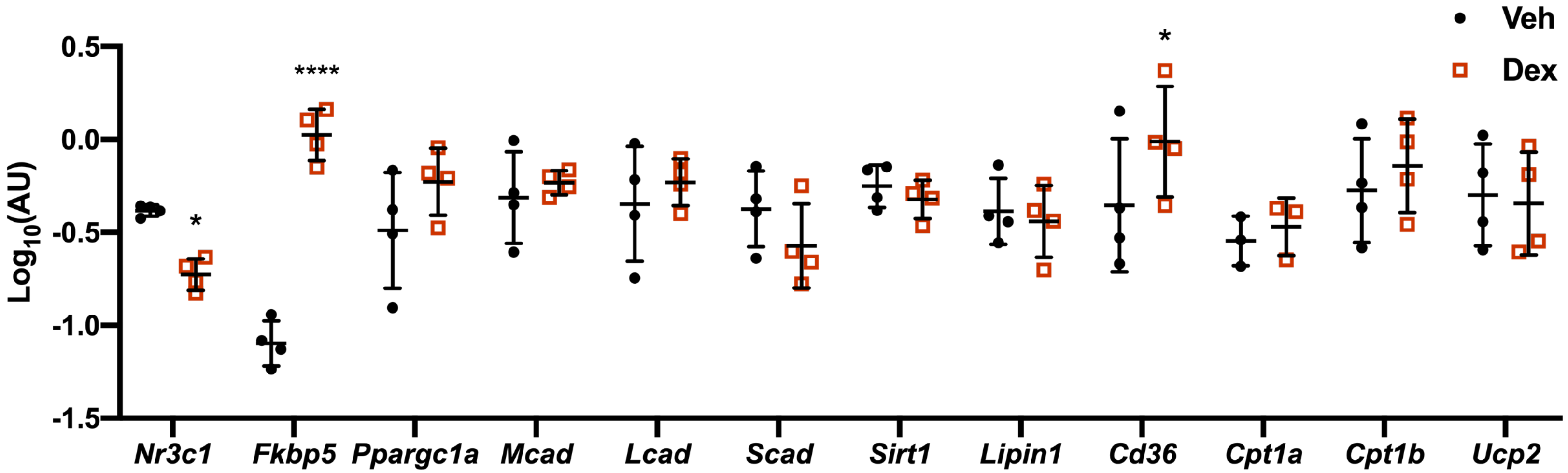


***SUPPLEMENTARY FIGURE 3.*** **Dexamethasone decreases GR expression but increases FKBP5 and CD36 expression in primary cultures of fetal cardiomyocytes, 6h after addition**

Primary fetal cardiomyocytes were prepared by digesting pooled E14.5-15.5 C57BL/6J fetal hearts then cultured under standard conditions for 48h before treatment with dexamethasone (Dex, 1μM) or vehicle (Veh, 0.01% ethanol) for 6h. Cardiomyocytes were then lysed in TRIzol and RNA isolated for analysis by qRT-PCR. Data are mean ± SD and were analysed by two-way ANOVA followed by post hoc Sidak's tests where *$p<0.05$, ****$p<0.0001$, n=4 independent cardiomyocyte pools, prepared on different days.

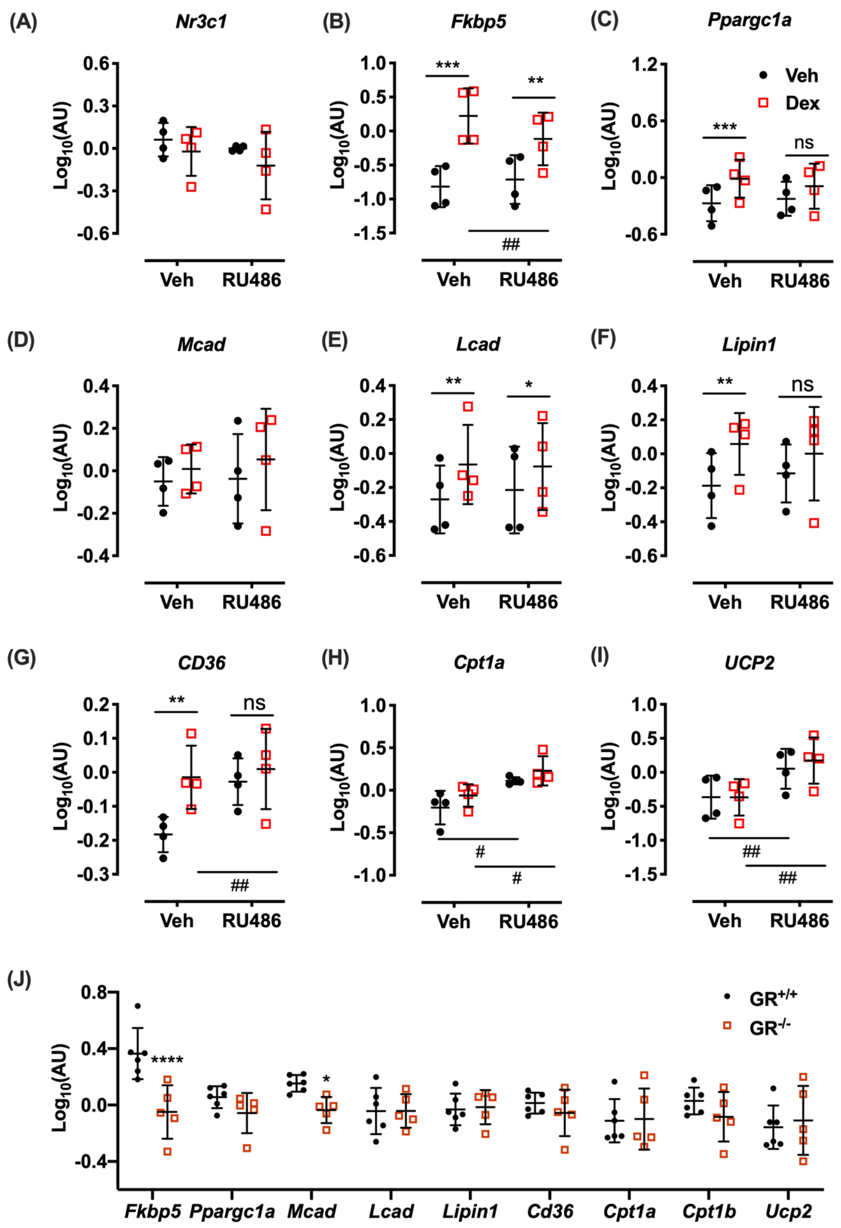


***SUPPLEMENTARY FIGURE 4.*** **GR antagonism attenuates the dexamethasone-induction of genes involved in mitochondrial fatty acid oxidation and some of these changes are seen in GR$^{-/-}$ fetal hearts at E17.5**

(A-I) Primary fetal cardiomyocytes were prepared by digesting pooled E14.5-15.5 C57BL/6J fetal hearts then cultured for 48h under standard conditions before being treated with the GR antagonist, RU486 (1μM) or vehicle (control, 0.01% ethanol) 30min prior to treatment with dexamethasone (Dex, 1μM) or vehicle (Veh, 0.01% ethanol). After 24h, cardiomyocytes were lysed in TRIzol and RNA isolated for analysis by qRT-PCR. (J) RNA was extracted from hearts of E17.5 fetal GR$^{+/+}$ and GR$^{-/-}$ mice for analysis by qRT-PCR. Data are mean ± SD, *p<0.05, ***p<0.01, **p<0.001 for comparisons between Veh/Dex, # p<0.05, ##p<0.01 for comparisons between control and RU486 treated cells by two-way ANOVA followed by post hoc Sidak's tests, n=4 independent cardiomyocyte pools, prepared on different days (A-I); n=5-6 individual fetuses from 5 different litters (J).

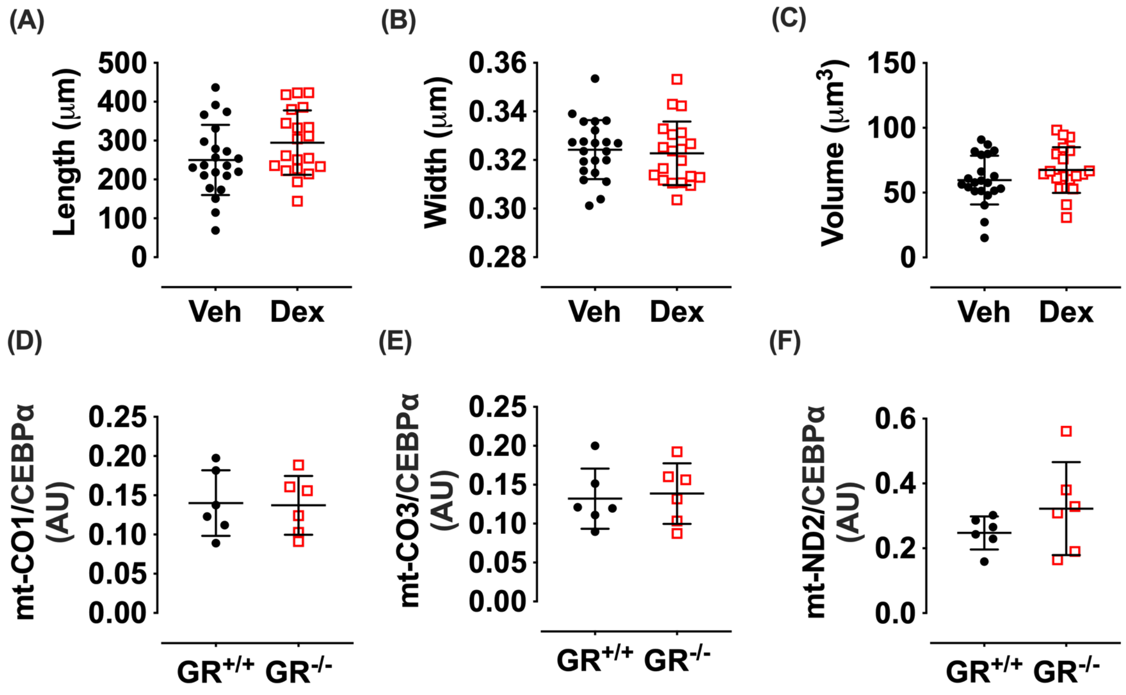


***SUPPLEMENTARY FIGURE 5.*** **Dexamethasone does not alter mitochondrial morphology nor does mitochondrial DNA content differ between $GR^{+/+}$ and $GR^{-/-}$ fetal hearts**

Primary fetal cardiomyocytes were isolated from pooled E14.5-15.5 fetal hearts from *mito*-QC (A-C) or C57Bl/6J mice (D-F). (A-C) Cardiomyocytes were cultured under standard conditions for 48h, then treated with dexamethasone (Dex, 1μM) or vehicle (Veh, 0.01% ethanol) for 24h and fixed with 4% PFA. Z-stacks of individual cells were obtained using a spinning disk confocal microscope. Mitochondrial morphology, including (A) length, (B) width and (C) total volume, was assessed using MitoGraph software (24, 25). (D-F) Mitochondrial DNA content of E17.5 fetal hearts from $GR^{+/+}$ and $GR^{-/-}$ littermates was quantified by qPCR of mitochondrially encoded genes: cytochrome c oxidase I (*mt-CO1*, **D**), III (*mt-CO3*, **E**) and NADH dehydrogenase II (*mt-ND2*, **F**), relative to the nuclear encoded (and intronless) *Cebpa* gene. Data are mean ± SD and were analysed by Student's t-tests: n=19-23 individual cells from a single pool of cardiomyocytes seeded in triplicate wells (A-C), n=6 fetuses from 5 litters (D-F).
